# Neuralized regulates a travelling wave of Epithelium-to-Neural Stem Cell morphogenesis in *Drosophila*

**DOI:** 10.1101/2020.06.02.129353

**Authors:** Chloé Shard, Juan Luna-Escalante, François Schweisguth

**Affiliations:** Institut Pasteur, Paris, France; UMR3738, CNRS, Paris, France; Laboratoire de Physique, Ecole Normale Supérieure, CNRS, Sorbonne Université, Université Paris Diderot, Paris, France

## Abstract

Many tissues are produced during development by specialized progenitor cells emanating from epithelia via an Epithelial-to-Mesenchymal Transition (EMT). Most studies have so far focused on cases involving single or isolated groups of cells. Here we describe an EMT-like process that requires tissue level coordination. This EMT-like process occurs along a continuous front in the *Drosophila* optic lobe neuroepithelium to produce neural stem cells (NSCs). We find that emerging NSCs remain epithelial and apically constrict before dividing asymmetrically to produce neurons. Apical constriction is associated with contractile myosin pulses and requires the E3 ubiquitin ligase Neuralized and RhoGEF3. Neuralized down-regulates the apical protein Crumbs via its interaction with Stardust. Disrupting the regulation of Crumbs by Neuralized led to defects in apical constriction and junctional myosin accumulation, and to imprecision in the integration of emerging NSCs into the transition front. Neuralized therefore appears to mechanically couple NSC fate acquisition with cell-cell rearrangement to promote smooth progression of the differentiation front.

## Introduction

During animal development, many complex organs are built by stem and progenitor cells originating from epithelial tissues (Norden, 2017). For instance, in vertebrate embryos, the anterior brain is produced by neuroepithelial cells that function as Neural Stem Cells (NSCs) and several organs, such as the intestine and the pancreas, derive from endodermal epithelia. Acquisition of stem-like properties has been associated with Epithelial-to-Mesenchymal Transition (EMT), at least in the context of carcinoma (Dongre and Weinberg, 2019). Thus, understanding how stem cells emerge in space and time from epithelia to build organs is an important issue.

During *Drosophila* neurogenesis a fixed number of NSCs, also termed neuroblasts, emerge from within the embryonic neuroectoderm via an Epithelial-To-Mesenchymal (EMT)-like delamination process to build the larval brain (Hartenstein and Campos-Ortega, 1984; Hartenstein and Wodarz, 2013). Each NSC has a unique spatial identity that is provided in part by the segmentation genes (Urbach and Technau, 2003). As neuroectodermal cells become specified as NSCs, they undergo apical constriction driven by a pulsatile actomyosin meshwork (Simões et al., 2017; An et al., 2017; Sawyer et al., 2010; Martin and Goldstein, 2014). Concomitantly, Adherens Junctions (AJ) are disassembled as E-cadherin (E-cad) is removed from the apical-lateral membrane, leading to NSC ingression (Simões et al., 2017). Delaminating NSCs, however, maintain apical-basal polarity cues such that, upon division, cell fate determinants localize at the basal pole and the mitotic spindle lines up along this polarity axis (Schober et al., 1999). As a result, NSCs divide asymmetrically to both self-renew and produce differentiated cells, neurons and glia (Knoblich, 2008).

A slightly different EMT-like process occurs in the larval brain for the formation of the Optic Lobe (OL) NSCs (Fig.1A). The outer proliferation center of the OL comprises a single layer of pseudostratified neuroepithelium (NE) cells that arise during embryogenesis and grow during larval development (Hofbauer and Campos-Ortega, 1990; Egger et al., 2007; Hakes et al., 2018). Then, in response to a systemic pulse of ecdysone, a self-perpetuating proneural wave sweeps the NE from medial to lateral during the third instar, eventually converting all NE cells into NSCs by the end of larval development (Dillard et al., 2018; Yasugi et al., 2008). These NSCs give rise to the medulla neurons that will comprise part of the visual processing center of the adult brain. This proneural wave is proposed to travel through the NE via an excitable reaction-diffusion mechanism. Accordingly, an activator signal, the Epidermal Growth Factor (EGF) Spitz, spreads in the NE by diffusion and self-induction and triggers a negative feed-back via the production of a non-diffusible inhibitor, i.e. the proneural factor Lethal of scute (L’sc) that inhibits the production of active Spitz by promoting NE-to-NSC differentiation (Jörg et al., 2019). This creates a traveling front of EGFR signaling and proneural activity (Fig.1B). The traveling domain of proneural gene expression is called the transition zone (TZ). The TZ is therefore defined as a stripe of 3-4 epithelial cells expressing the proneural factor L’sc. Spitz is secreted by medial TZ cells and diffuses to activate EGF Receptor (EGFR) signaling in more lateral NE, whereas L’sc induced by EGFR both self-activates and regulates the NE-to-NSC fate transition via Delta-Notch signaling (Wang et al., 2011; Weng et al., 2012; Yasugi et al., 2008, 2010; Sato et al., 2016; Jörg et al., 2019). NE cells therefore appear to follow a stereotyped fate trajectory as the proneural wave sweeps the NE. Starting from a Notch^ON^ / proneural^OFF^ state in the NE, cells entering the TZ express low but increasing levels of L’sc to acquire a Notch^ON^ / proneural^ON^ state before switching to a Notch^OFF^ / proneural^ON^ state as they adopt the NSC fate (Egger et al., 2010; Weng et al., 2012; Ngo et al., 2010; Wang et al., 2011; Orihara-Ono et al., 2011). The ON/OFF switch in Notch activity likely results from high levels of Delta, cis-inhibiting Notch in the TZ cells located close to the NSCs (Del Álamo et al., 2011; Contreras et al., 2018; Reddy et al., 2010). While the logic underlying progression of the TZ and cell fate switch within the TZ is partly understood, how epithelial NE cells undergo cell shape and polarity changes to become non-epithelial NSCs remains unexplored. Interestingly, while this epithelium-to-NSC transition exhibits similarities with NSC delamination in the embryo, there are also notable differences. First, instead of occurring in individual cells as seen in the embryonic neuroectoderm, this EMT-like transition involves a continuous stripe of cells undergoing epithelium remodeling in a coordinated manner at the tissue level. Second, newly-specified NSCs do not delaminate from the NE to locate basally but rather remain in the same plane as NE cells (Ngo et al., 2010; Egger et al., 2007; Yasugi et al., 2008). Thus, the OL NE provides an opportunity to examine how an EMT-like process producing NSCs is regulated and coordinated at the tissue level.

**Figure 1:**
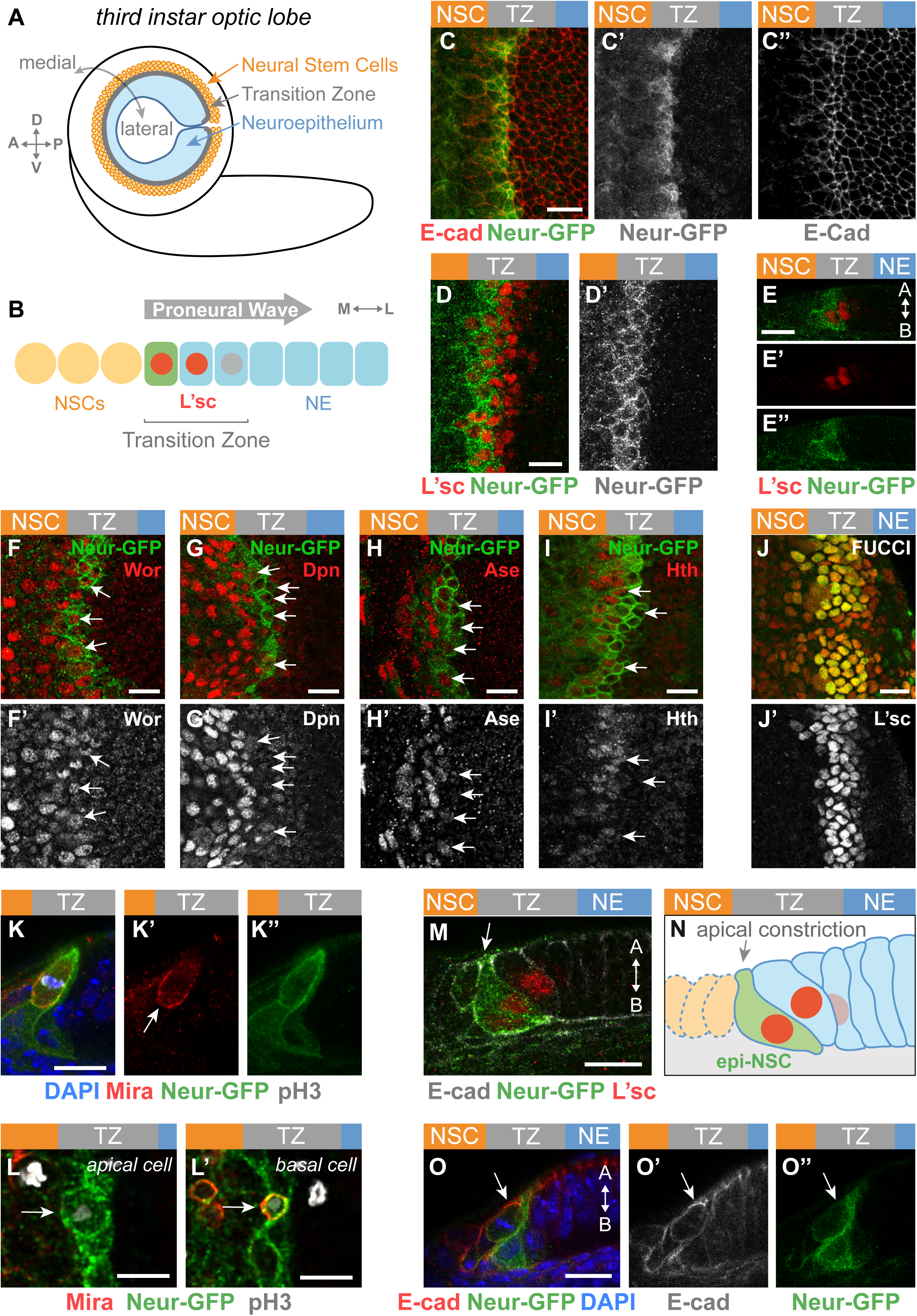
Neur marks the asymmetrically-dividing TZ cells producing the first-born neurons. **(A)** Diagram of the third instar larval brain displaying the OL neuroepithelium (NE), transition zone (TZ) and medulla neural stem cells (NSCs). P, posterior, A, anterior, D, dorsal, V, ventral. **(B)** The proneural wave sweeps the NE from medial to lateral inducing NE-NSC transition in a region called the TZ defined by the graded expression of L’sc (red nuclei). The medial TZ cell appears in green (M, medial, L, lateral). (**C-E’’**) Surface (C-D’) and cross-section views (E-E’’; A, apical, B, basal) showing that Neur-GFP (green) was detected in the medial-most TZ cells (L’sc, red in D-E’’). These cells showed increased E-cad levels (red in C). Lower levels of Neur-GFP were observed in NSCs (C’,D’,E’’). (**F-I’**) All Neur-positive TZ cells (Neur-GFP, green) co-expressed Wor (red, F) while Neur expression partially overlapped with the NSC markers Dpn (red, G), Ase (red, H) and Hth (red, I). Arrows point to the nucleus of several medial TZ cells. (**J**,**J’**) TZ cells (L’sc, J’) expressed CycB-RFP (red, J) and E2F1-GFP (green, J). All TZ cells expressed both FUCCI markers, indicating that they are in G2 phase. (**K-L’**) Medial TZ cells (Neur-GFP, green) appeared to undergo the first round of asymmetric cell division (pH3, phospho-histone H3 in white; DAPI, blue). This division was oriented along the apical-basal axis and Mira localized asymmetrically (K-K’’; Mira, red). The arrow in K’ indicates the basal crescent of Mira. A large apical cell (L) and a small basal cell (L’), likely corresponding to the first-born GMC, appeared to be produced by this asymmetric division. (**M-O’’**) Cross-section views showing that the medial-most TZ cells marked by Neur-GFP (green) and L’sc (red in M) were apically constricted and had a characteristic ‘tear drop’ shape while retaining apical Adherens Junctions (M; E-cad, white). E-cad remained detectable as these cells divide (O-O’’; E-cad, red; DAPI, blue). Since these apically-constricted epithelial cells underwent the first round of asymmetric cell divisions, they were named epi-NSCs (N). Scale bars = 10 μm.

The protein Crumbs (Crb) is a large transmembrane protein with a short intracellular tail that recruits the scaffolding protein Stardust (Sdt in flies, Pals1 in mammals). Sdt interacts with Patj to form a Crb-Sdt-Patj complex (Bulgakova and Knust, 2009). This Crb complex is required to stabilize AJs and maintain epithelial integrity during tissue remodeling in *Drosophila* (Harris and Tepass, 2008; Bajur et al., 2019; Flores-Benitez and Knust, 2015; Campbell et al., 2009; Perez-Mockus et al., 2017b; Röper, 2012; Salis et al., 2017). Recently, the conserved E3 ubiquitin ligase Neuralized (Neur) was found to directly interact with specific isoforms of Sdt and this interaction appeared to target Crb for endocytosis and degradation (Perez-Mockus et al., 2017b). This activity of Neur was found to promote epithelium remodeling in the embryonic gut and increased Neur activity in the embryonic ectoderm resulted in a loss of epithelial integrity (Chanet and Schweisguth, 2012; Perez-Mockus et al., 2017b). These findings led to suggest that Neur expression in NSCs might promote their EMT-like delamination (Perez-Mockus and Schweisguth, 2017). Whether Neur regulates the epithelium-to-NSC transition in the OL is not known.

Here we studied how TZ cells undergo an epithelium-to-NSC transition. We found that TZ cells adopting a NSC fate apically constrict one cell at a time and remain epithelial until they divide asymmetrically to produce the first-born differentiated cells. Neur becomes expressed in these newly-specified NSCs and regulates apical constriction in part by down-regulating Crb in a Sdt-dependent manner. The resulting anisotropy of Crb accumulation appeared to promote the accumulation of junctional Myosin II (MyoII) and formation of supra-cellular actomyosin cables within the TZ. Live imaging of epithelium-to-NSC transition suggests a model whereby regulation of Crb by Neur in the TZ contributes to coordinate fate acquisition at the single cell level with epithelium remodeling at the tissue level to ensure a smooth progression of the differentiation front.

## Results

### Emerging NSCs retain epithelial properties

Previous studies have indicated that TZ cells switch off Notch signaling as they become NSCs (Egger et al., 2010; Weng et al., 2012; Ngo et al., 2010; Wang et al., 2011; Orihara-Ono et al., 2011) (Fig.1B). By analogy to the neural progenitors that switch off Notch and express Neur as they are singled-out from proneural clusters (Huang et al., 1991; Castro et al., 2005; Rouault et al., 2010), Neur could be a marker of the TZ cells that switch off Notch signaling. Recently, Neur was proposed to act in the medial-most TZ cells to up-regulate the endocytosis of Delta, thereby trans-activating Notch in neighboring TZ cells (Contreras et al., 2018; Weinmaster and Fischer, 2011). Previous analysis of Neur expression in the OL relied on a *neur-lacZ* transcriptional reporter (Contreras et al., 2018). Here, to better characterize where Neur is expressed, we used a functional BAC transgene expressing Neur-GFP under the control of the *neur* genomic regulatory sequences (Perez-Mockus et al., 2017a). We found that Neur accumulated in the TZ cells that are located close to the NSCs (Fig.1C-C’’). Notably, Neur had a more restricted domain of expression than L’sc, although their expression overlapped Neur was expressed in approximately one row of cells at the medial edge of the NE (Fig.1D-E’’). It was also detected at lower levels in NSCs. Moreover, all Neur-positive TZ cells expressed Worniu (Wor), a transcription factor associated with NSC identity (Ashraf and Ip, 2001; Cai et al., 2001) (Fig. 1F,F’), whereas Deadpan (Dpn), Asense (Ase) and the temporal identity factor Homothorax (Hth) (Suzuki et al., 2013; Li et al., 2013) displayed only partial overlap with Neur-GFP (Fig.1G-I’). Thus, the Neur-positive TZ cells have a transcriptional identity that is an intermediate between NE and NSC fates and Wor is an early NSC identity factor during the NE-NSC transition.

We next studied the division mode of the Neur-positive TZ cells. As NE cells differentiate into NSCs, they gain the ability to undergo asymmetric division to produce another NSC and a neuron-producing Ganglion Mother Cell (GMC) (Egger et al., 2007; Hofbauer and Campos-Ortega, 1990). Given that Neur-positive TZ cells express NSC identity genes, we wondered whether they divide asymmetrically like NSCs. While earlier observations suggested that TZ cells are arrested in the G1 phase of the cell cycle (Weng et al., 2012; Orihara-Ono et al., 2011; Reddy et al., 2010), we first found using the FUCCI (fluorescent ubiquitination-based cell cycle indicator) system (Zielke et al., 2014) that all TZ cells are in G2 (Fig.1J,J’). Consistent with this, Cyclin B was found to accumulate in TZ cells (Fig.S1). Furthermore, we observed dividing Neur-positive TZ cells with asymmetric Miranda (Mira) (Fig.1K,K’) (Ikeshima-Kataoka et al., 1997). Analysis of these cells at telophase/cytokinesis indicated that two cells of different sizes were produced from the division (Fig.1L,L’). The smaller daughter cell located basally appeared to be a GMC, based on Prospero (Pros) staining (Spana and Doe, 1995) (Fig.1L’, Fig.S1). Given that Neur cells co-express Hth, this GMC is likely to be the first Hth lineage born. Together, these data indicate that TZ cells are in G2 and that medial TZ cells undergo the first round of asymmetric division to produce early-born GMCs.

It was previously reported that the medial-most TZ cells have a characteristic teardrop shape suggestive of apical constriction (Orihara-Ono et al., 2011). We confirmed this observation by showing that the Neur-positive TZ cells apically constrict (Fig.1M-N). Importantly, these TZ cells remain epithelial with apical E-cad, and retain E-Cad as they undergo mitotic rounding and asymmetric division (Fig.1O).

Finally, we examined the division of these Neur-positive TZ cells using *ex vivo* live imaging of L3 larval brains expressing Neur-GFP and Histone2Av-RFP (Fig.S1). Tracking cells live, we observed that medial Neur-positive TZ cells divide asymmetrically to produce a small basal daughter cell with high Neur levels. Live imaging of larval brains expressing E-cad-Cherry and Neur-GFP further revealed that medial Neur-positive TZ cells exhibit a clear E-cad signal as they round up during mitosis, indicating that these TZ cells retain apical junctions when they enter mitosis (Fig.S1).

In summary, our data strongly suggest that the medial TZ cells are epithelial cells that undergo a first round of asymmetric cell division to produce the earliest-born GMCs. Thus, the medial Neur-positive cells of the TZ have unique features: they express early NSC markers and undergo the first round of asymmetric cell division but differ from more mature NSCs by the presence of apical E-cad, a key feature of epithelial cells that is lost in NSCs (Fig.1O; see also below). We refer here to this transitory epithelial-like NCS state as the epi-NSC state and to the Neur-positive TZ cells as the epi-NSCs.

### Apical constriction by medial Myosin in epi-NSCs

While epi-NSCs retain epithelial features until they divide, they appear to undergo apical constriction which is suggestive of changes in epithelial organization. To better characterize this morphogenetic process, we used E-cad staining to segment cells and quantify apical area (Fig.2A-B). This analysis showed that medial TZ cells are indeed apically constricted. We further observed that apical constriction resulted in a slightly higher density of epi-NSCs compared to other TZ cells (Fig.2C), suggestive of cell-cell intercalation events within the TZ. In many epithelia, apical constriction can be induced by a contractile actomyosin meshwork (Martin et al., 2009; Martin and Goldstein, 2014). To monitor and correlate myosin dynamics with changes in apical area, we performed live imaging on *ex vivo* cultured brains from L3 larvae expressing a GFP-tagged allele of Myosin II (MyoII-GFP) and E-cad-mCherry (or Par3-mCherry) to mark apical junctions. In these movies, epi-NSCs were detected as a row of apically constricted cells. Individual epi-NSCs undergoing a decrease of their apical area prior to division were selected for analysis. We observed that these epi-NSCs apically constricted through phases of apical area contraction and expansion (Fig.2D-F and movie S1).

**Figure 2:**
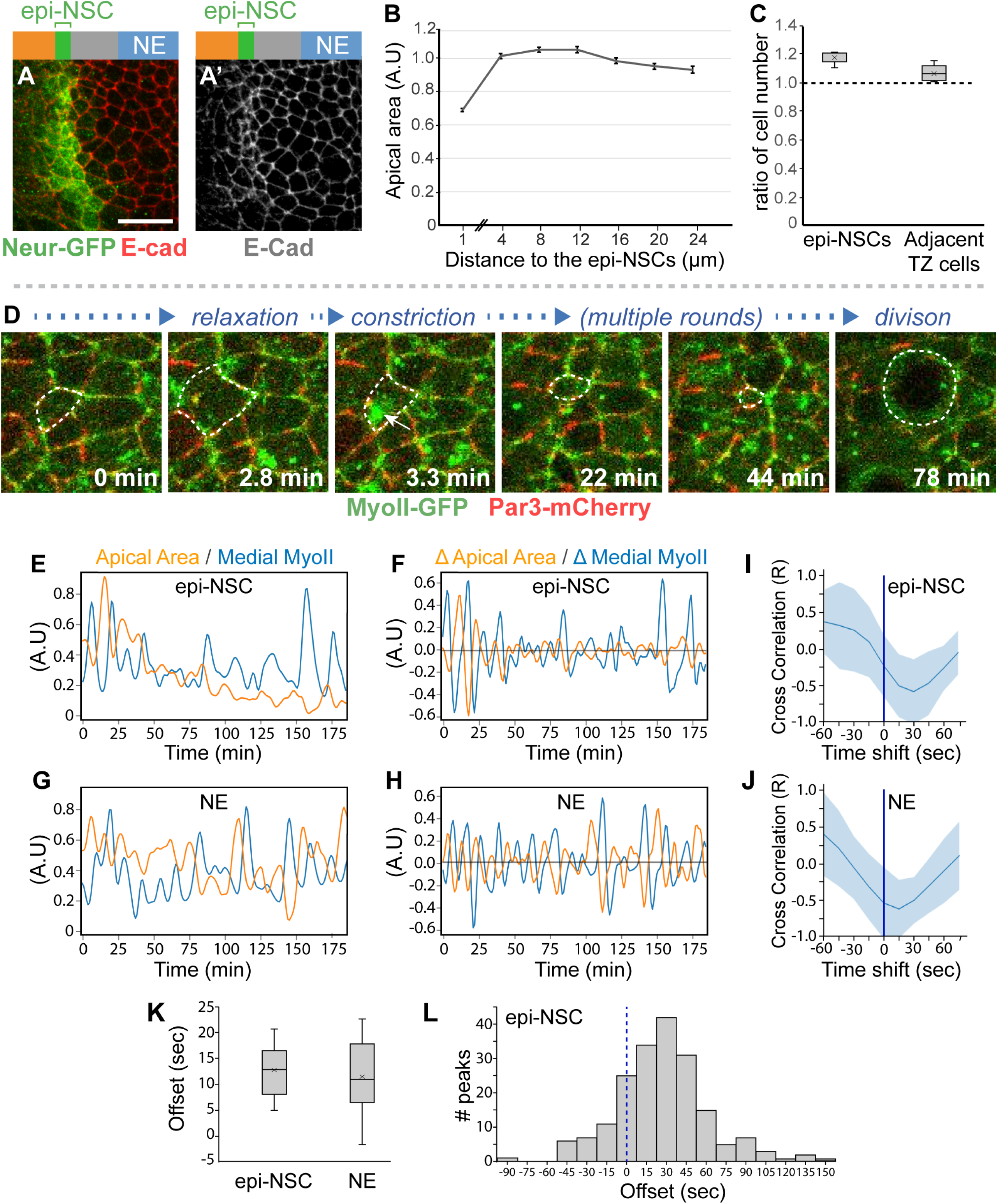
Apical constriction of epi-NSCs correlates with pulses of apical MyoII levels. **(A)** Surface view of wild-type NE stained for E-cad (red) and Neur-GFP (green) showing that epi-NSCs have smaller apical areas. **(B)** Quantification of apical area (normalized, A.U., arbitrary units) plotted against distance from epi-NSCs (in μm; x=0 corresponds to a line joining the center of each epi-NSCs) showed that apical area decreases towards the medial edge of the TZ (*n* = 13 brains; means ± SEM of all pooled cells, 373-890 cell per binned distance). **(C)** Density of epi-NSCs and other TZ cells along the D-V axis relative to those of lateral NE cells (*n* = 8 brains; boxplots display the mean ratios, with the density of distal NE cells normalized to 1 for each brain). **(D)** Snapshots from a movie of a brain expressing MyoII-GFP (green) and Par3-mCherry (red) showing an apically constricting TZ cell (dotted outline) undergoing phases of constriction and expansion. Constriction is associated with increased levels of medial-apical MyoII (arrow). (**E-H**) plots of apical area and MyoII intensity values over time for a representative epi-NSC and NE (G) cells, along with the rates of changes in apical area and MyoII levels (F,H). Increased MyoII levels was associated with reduced apical area in both epi-NSC and NE cells. (**I**,**J**) Cross-correlation values between area and myosin rates of change (mean +/-SEM calculated for the different peaks) plotted as a function of the time-shift in representative epi-NSC (**I**) and NE cells (**J**). Anti-correlation was maximal with a 10-20s time-shift, i.e. offset, with maximal increase of MyoII preceding maximal rate of apical constriction. **(K)** Mean offset time between maximal rates of MyoII accumulation and area changes in epi-NSCs (*n* = 9) and NE cells (*n* = 10). MyoII accumulation similarly precedes constriction in both cell types (no significant difference, two-tailed *t* test). **(L)** Offset times between maximal accumulation of MyoII and maximal constriction rates in epi-NSCs (*n* = 191 peaks, from 9 NSCs).

Additionally, the levels of apical-medial MyoII appeared to pulse, suggesting phases of recruitment of MyoII at, and dissociation from, the apical cortex (Fig.2D-F). These MyoII pulses negatively correlated with apical area changes (Fig.2I). In addition, maximal accumulation of medial MyoII appeared to precede apical constriction by approximately 13 +/-8 seconds (sec) on average in all epi-NSCs analyzed (Fig.2K). In 46% of apical constriction phases, the closest maximal accumulation of medial MyoII in time preceded apical reduction within 45 sec (Fig.2L). Together, these data suggest that pulses of apical-medial MyoII drive apical constriction in epi-NSCs, as previously observed in several other epithelia (Martin et al., 2009; Martin and Goldstein, 2014), including in the embryonic neuroectoderm during NSC delamination (Simões et al., 2017; An et al., 2017).

We next compared MyoII dynamics in epi-NSCs with those of the NE cells that were not undergoing apical constriction. Both reduction of the apical area and medial MyoII pulses appeared to transiently occur in the NE (Fig.2G-J and movie S2) with maximal accumulation of medial MyoII preceding constriction by 12 +/-11 sec (Fig.2K). This indicated that pulses of apical-medial MyoII triggered constriction in both NE and epi-NSCs. However, in contrast to epi-NSCs, NE cells did not apically constrict over time but rather displayed balanced phases of contraction and expansion. Thus, considering that apical constriction results from a pulse of medial actomyosin that persists long enough to let the energy stored dissipate in part via the remodeling of apical junctions and associated actin cortex (Martin et al., 2009; Martin and Wieschaus, 2010), we suggest that epi-NSCs might differ from NE cells in cortex remodeling such that remodeling is fast enough in epi-NSCs, but not in NE cells, to stabilize a loss of apical area upon constriction (Clement et al., 2017). In conclusion, contractile MyoII pulses appeared to direct sustained apical constriction in epi-NSCs but not in NE cells.

### Junctional MyoII accumulation suggestive of supra-cellular cables within the TZ

While studying MyoII dynamics, we observed high levels of junctional MyoII at cell-cell contacts between epi-NSCs and the more lateral TZ cells (Fig.3A-A’’). Increased junctional MyoII was particularly noticeable where epi-NSCs line up to create a straight tissue interface with other TZ cells (Fig.3A’’). We quantified the intensity of junctional MyoII along cell-cell contacts parallel to the differentiation front at four positions: at the NSC/epi-NSC and epi-NSC/TZ boundaries (a and b junctions, respectively), at the next cell-cell contacts within the TZ (c junctions) and within the NE (d junctions) (Fig.3B). This quantification showed that the epi-NSC/TZ junctions had significantly increased MyoII levels (Fig.3C). Additionally, increased junctional MyoII was accompanied by elevated accumulation of F-actin (Fig.3H-H’’’). These observations suggested that supra-cellular cables of actomyosin may form to produce enough tensile forces to line up epi-NSCs along this tissue interface (Roper, 2013). However, given the architecture of the brain, with the optic lobe epithelium sandwiched between glial cells at the brain surface and differentiating neural tissues underneath, probing forces is technically challenging and was not performed.

**Figure 3:**
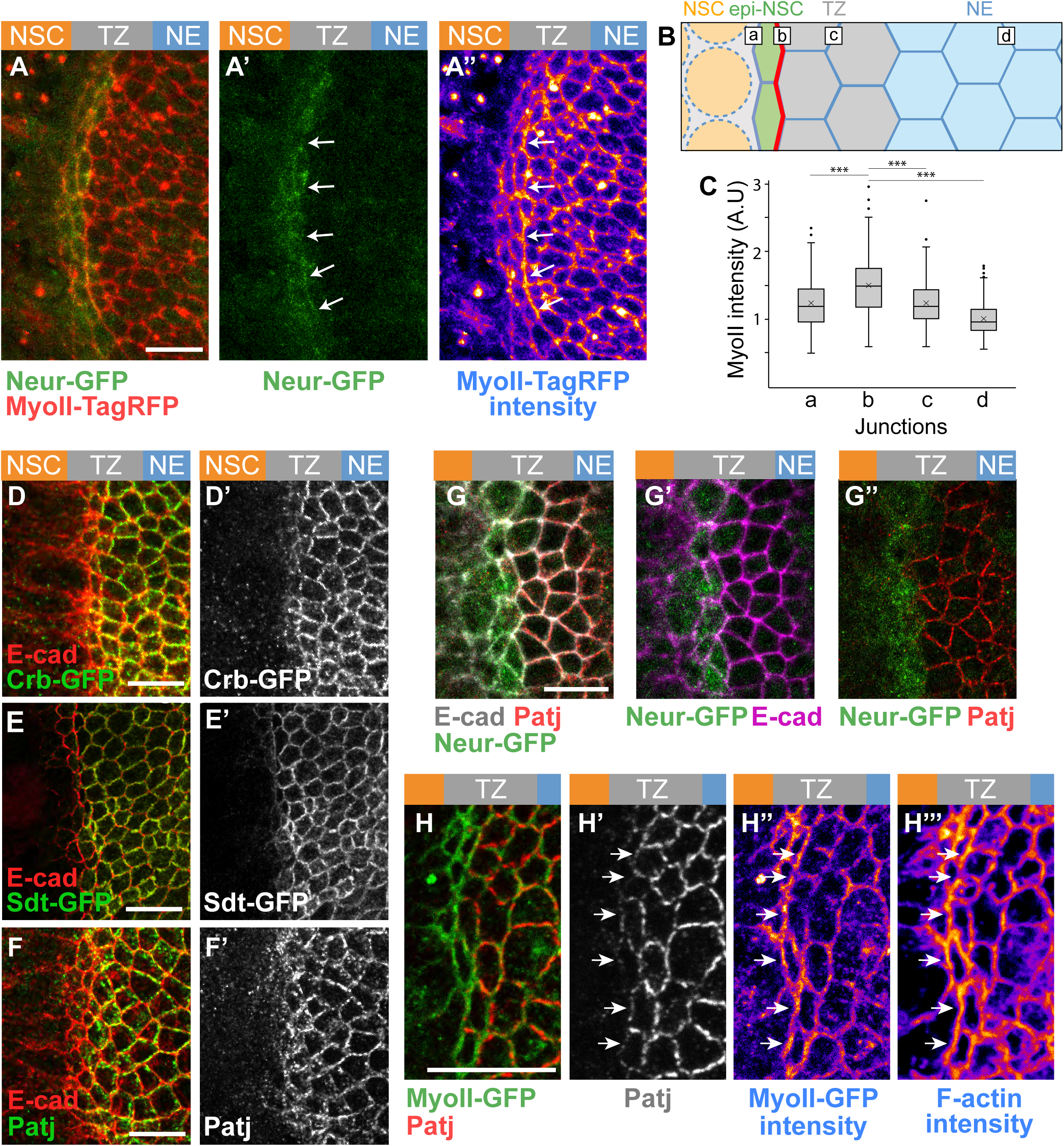
Low Crb in epi-NSCs correlates with polarized junctional MyoII. (**A-A’’**) Snapshot of a brain expressing MyoII-TagRFP (red in A, intensity levels in A’’) and Neur-GFP (green) showing an enrichment of junctional MyoII at the cell-cell contact between epi-NSC and more lateral TZ cells (arrows). Neur localized at apical junctions (A’). **(B)** Diagram illustrating the different cell-cell contact used for the quantification of junctional MyoII, i.e. between NSCs and epi-NSCs (a), epi-NSCs and immediately adjacent TZ cells (b), between Neur-negative TZ cells (c) and lateral NE cells (d). **(C)** Box-plots of junctional MyoII intensity at four distinct junctions as in (B). MyoII is significantly enriched at b junctions relative to all others (*n* = 6, p-values <10^−15^, one-way ANOVA, Scheffé multiple comparison tests). (**D-G’’**) Crb-GFP (green; D,D’), Sdt-GFP (green, all isoforms; E,E’) and Patj (green in F,F’; red in G-G’’), but not E-cad (red in E,F,G; magenta in G’), were specifically down-regulated in apically-constricted epi-NSCs (Neur-GFP, green, in G-G’’). (**H-H’’’**) MyoII-GFP (green, H) was enriched along junctions lacking Patj (red; see arrows in H’,H’’). F-actin (phalloidin staining, H’’’) was also enriched along junctions lacking Patj. Scale bars = 10 μm.

In other epithelia, supra-cellular MyoII cables have been observed along boundaries of cells with different levels of the apical protein Crb, and an anisotropic loss of Crb can cause their formation (Röper, 2012). To test whether Crb differentially accumulates in epi-NSCs relative to other TZ cells, we studied the pattern of Crb accumulation using GFP-tagged Crb. We found that Crb localized at apical junctions in NE and lateral TZ cells but was lost in medial epi-NSCs (Fig.3D). The two other core components of the Crb complex, Sdt and Patj, were similarly down-regulated in epi-NSCs (Fig.3E-G’’). In contrast, the levels of E-cad and Par3 at apical junctions appeared largely unchanged (Fig.S2; note, however, that E-cad levels appeared to be slightly higher whereas Par3 was in part cytoplasmic in apically-constricted epi-NSCs, suggestive of junction remodeling). Additionally, co-staining for Patj with MyoII-GFP showed that MyoII was enriched at junctions where Patj was downregulated (Fig.3H-H’’’). These observations support a model whereby low Crb allows for junctional remodeling in epi-NSCs. In summary, our data suggest that supra-cellular actomyosin cables form at junctions juxtaposing non-apically constricted TZ cells with high Crb and apically constricted epi-NSCs with low Crb, possibly creating a tension interface within the OL epithelium.

### Neur down-regulates Crb via Sdt in epi-NSCs

We next looked for candidate regulators of Crb that might down-regulate Crb in epi-NSCs and first considered Neur. Indeed, while a key function of Neur is to target Delta for endocytosis (Lai et al., 2001; Le Borgne and Schweisguth, 2003; Weinmaster and Fischer, 2011), Neur is also known to down-regulate Crb, Std and Patj, possibly via endocytosis and degradation of Crb complexes (Fig. 4A) (Perez-Mockus et al., 2017b). Neur directly interacts with a specific subset of Sdt isoforms, i.e. those that contain a Neur Binding Motif encoded by exon 3, and thereby targets for degradation the Crb complexes that include these Sdt isoforms (Fig.4B). Using a knock-in allele of *sdt* that has GFP inserted into the sequence of exon 3 (Perez-Mockus et al., 2017b), we found that the Neur-regulated isoforms of Sdt are expressed in the NE and TZ (Fig.4C, E). We therefore tested the role of Neur in the down-regulation of Crb using *neur* mutant clones. Crb was found to co-localize with E-cad in *neur* mutant cells past the TZ boundary (Fig.4F-F’’). This indicates that Neur is required for the down-regulation of Crb complexes in epi-NSCs. Also, E-cad staining appeared to persist in NSCs close to the TZ (Fig.4F’’), suggesting a delay in AJ disassembly in mutant NSCs. Forced expression of the E3 ubiquitin ligase Mindbomb1 (Mib1), which can act redundantly with Neur in the regulation of Delta endocytosis (Lai et al., 2005; Le Borgne et al., 2005), did not rescue the delay in Patj down-regulation in *neur* mutant cells (Fig.S3). This indicates that Neur is unlikely to down-regulate Crb complex proteins in the OL via the regulated endocytosis of Delta. To test whether Neur down-regulates Crb via Sdt, we used a *sdt* allele that was engineered to flank the exon 3 with FRT sites such that only Neur-resistant isoforms of Sdt are expressed upon FLP-mediated excision of this exon (Fig.4C). Using this tool, we compared the localization of Crb complex proteins in the *optix-Gal4* or *hh-Gal4* cells expressing the Neur-resistant isoforms of Sdt with control cells in other domains of the NE (Fig.4G-G’’). Patj was maintained at the apical surface of the Neur-positive epi-NSCs when exon 3 of *sdt* is excised. Moreover, both Patj and E-cad persisted in NSCs marked by Ase or Dpn (Fig.4H-L’). We therefore conclude that Neur down-regulates the Crb complex via Sdt in epi-NSCs. Of note, we also observed an increase in Patj accumulation in lateral NE cells (Fig.S3). This suggests that the short Sdt isoforms produced upon deletion of exon 3 promote the stability of the Crb complex even in cells that do not express Neur.

**Figure 4:**
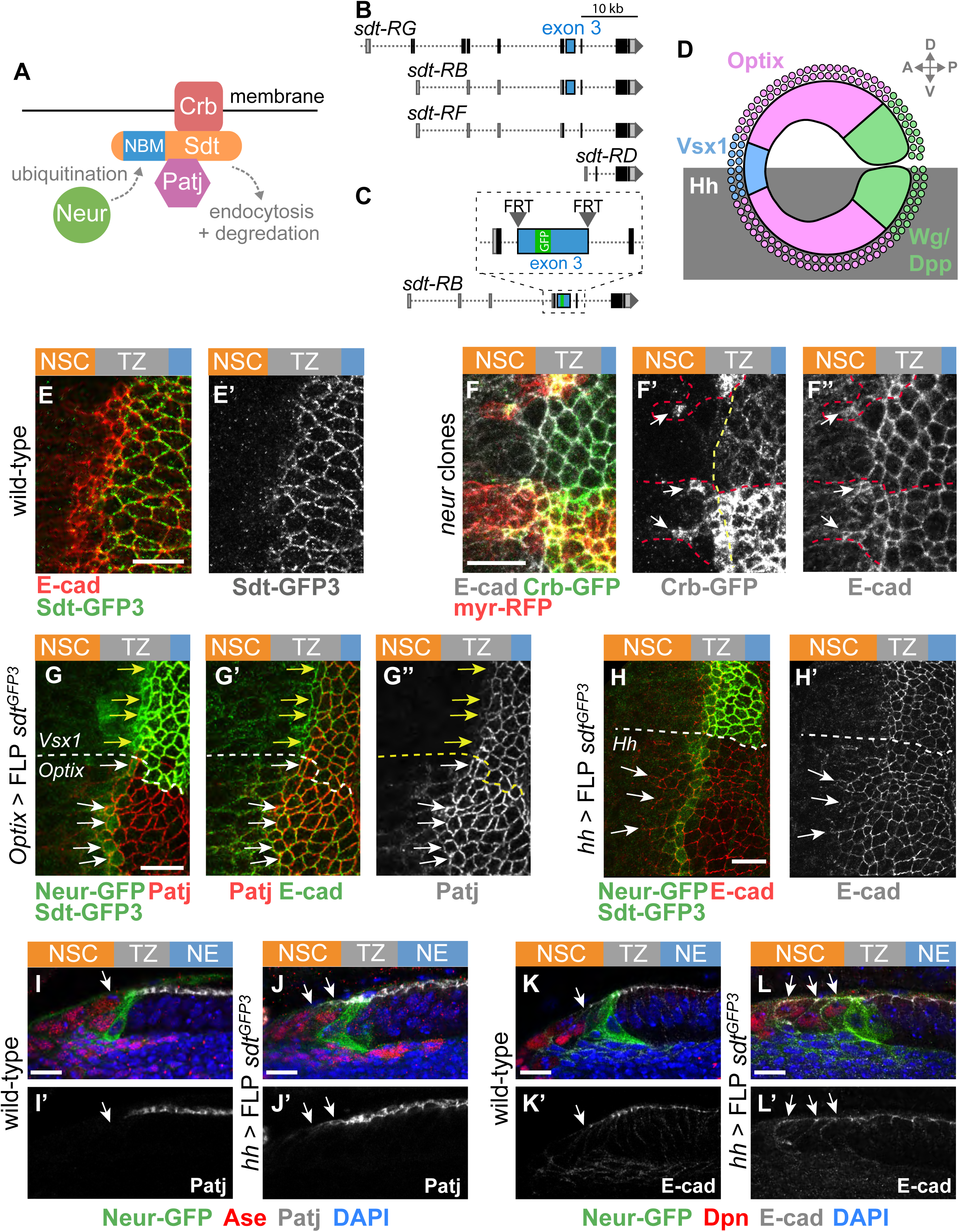
Neur acts in part via Sdt to down-regulate Crb in epi-NSCs. **(A)** Working model of Crb regulation by Neur. Neur interacts directly with Sdt isoforms that include the Neur Binding Motif (NBM) encoded by exon 3 and is proposed to target the Crb complex for endocytosis and degradation. **(B)** Simplified view of the alternatively spliced *sdt* transcripts. A subset of the *sdt* mRNAs, including *RB* and *RG* but not *RD* and *RF*, encodes exon 3. **(C)** Molecular structure of the *sdt*^*GFP3*^ allele with GFP inserted into exon 3, itself flanked by FRT sites. **(D)** Diagram of the NE subdivided into spatial domains expressing different transcription factors including Optix and Vsx1, or signaling molecules including Dpp, Wg, and Hh, hence providing a toolbox of spatially restricted Gal4s. (**E**,**E’**) Sdt isoforms containing the sequence encoded by exon3 (Sdt-GFP3, green) were expressed in the NE and down-regulated in epi-NSCs (E-cad, red). (**F-F’’**) Apical Crb (Crb-GFP, green) was detected colocalizing with E-cad (white) in *neur* mutant cells past the TZ medial edge (arrows in F’,F’’). MARCM *neur*^*IF65*^ clones were marked by the expression of a membrane RFP (myrRFP, red). (**G-H’**) Expression of only Neur-resistant isoforms of Sdt, via the excision of exon 3 in the *sdt*^*GFP3*^ allele using *optix-Gal4 UAS-FLP* led to the detection of Patj (red; white arrows in G-G’’) colocalizing with E-cad (green, G’) in mutant epi-NSCs (Neur-GFP, green; G) within the Optix domain of the NE (note the loss of Std-GFP3, green in G, in this domain). In contrast, Patj was down-regulated in control epi-NSCs of the Vsx1 domain (yellow arrows) which served as internal negative control (Sdt-GFP3, green; G). Excision of exon 3 of *sdt* in the Hh domain using *hh-Gal4 UAS-FLP* had a similar effect on Cad (red in H,H’; see loss of Std-GFP3, green in G, in this domain; Neur-GFP, also green). (**I-L’**) Cross-section views showing that Patj (white in I-J’) was down-regulated in epi-NSCs (Neur-GFP, green) in the presence of Neur-sensitive Sdt in wild-type brains (I,I’) but not when only Neur-resistant Sdt was expressed upon excision of exon 3 in *hh*>FLP *sdt*^*GFP3*^ (J,J’). Note that Patj was observed at the apical cortex of NSCs (Ase, red) upon excision of exon 3 (J,J’). Likewise, E-cad (white in K-L’) was detected further within the NSC domain (Dpn, red) when Neur resistant isoforms of Sdt were expressed (K-L’). DAPI (blue) marked all nuclei. Scale bars = 10 μm.

### Neur regulates apical constriction in epi-NSCs in part via Sdt

We next asked whether this regulation of Crb by Neur is required for the apical constriction of epi-NSCs. We found that medial TZ cells were no longer apically constricted in *neur* mutant clones (Fig.5A,A’). Additionally, ectopic expression of the Neur inhibitor Brd^R^, a stabilized version of Brd (Perez-Mockus et al., 2017a; Bardin and Schweisguth, 2006; De Renzis et al., 2006) where the lysine (K) residues have been mutated into arginine (R), strongly reduced apical constriction in addition to recapitulating the phenotype of apical E-cad in NSCs (Fig.5B’). Finally, we used RNAi against the *l’sc* gene to perturb the expression of the *neur* gene, a direct target of the proneural factors (Rouault et al., 2010; Miller et al., 2014; Contreras et al., 2018). We observed that knocking-down *l’sc* resulted in low and heterogeneous Neur-GFP expression (Fig.S4). Interestingly, we found that Neur-GFP intensity levels negatively correlated with apical area in epi-NSCs (Fig.S4). Together, these results show that Neur positively regulates the apical constriction of epi-NSCs.

**Figure 5:**
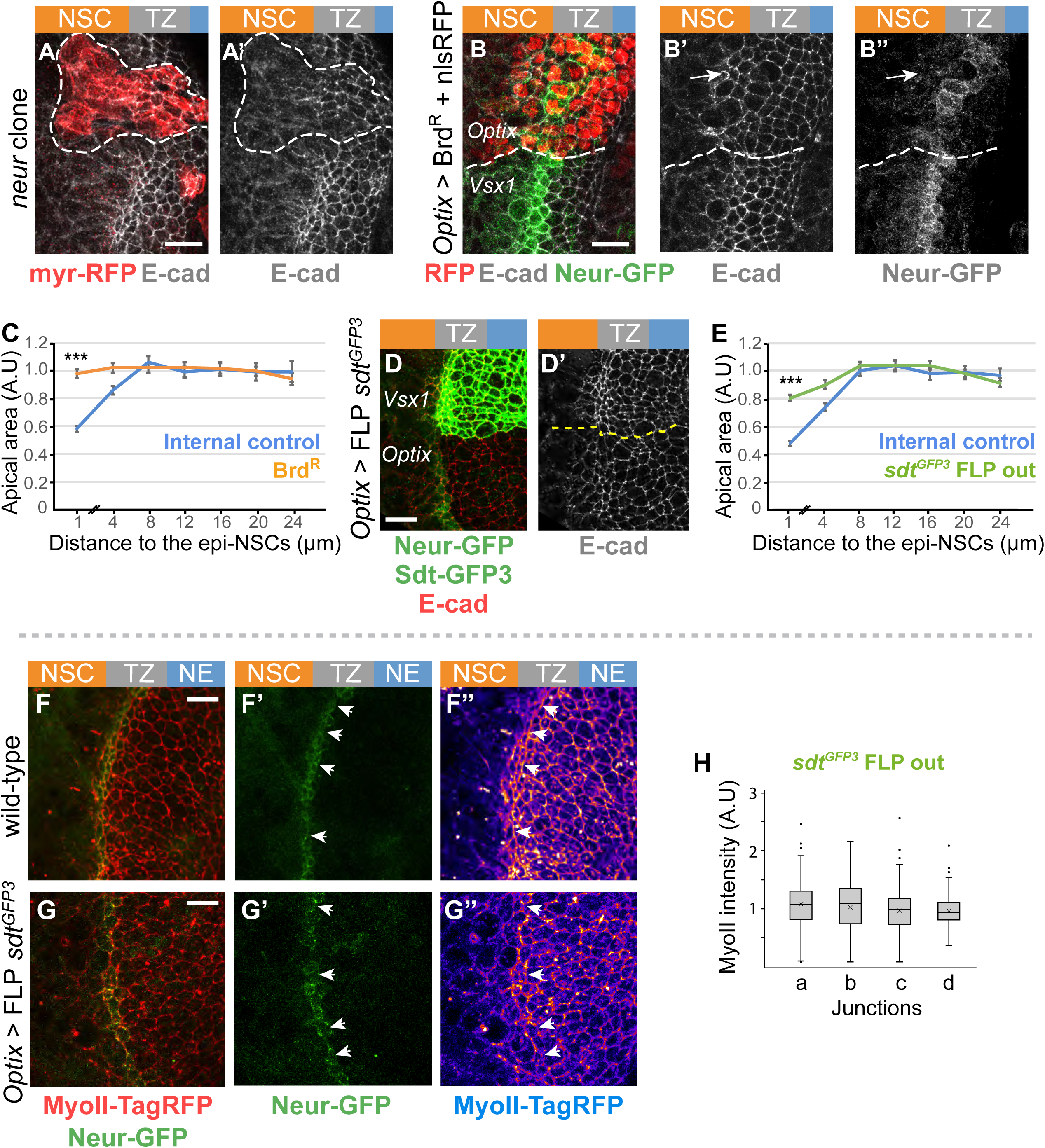
Neur is required for apical constriction and acts in part via Sdt. (**A**,**A’**) Loss of *neur* activity in *neur*^*IF65*^ MARCM clones (GFP, green) led to defective apical constriction in cells located at the medial edge of the TZ as well as to the persistence of E-cad (white) in mutant NSCs. (**B-B’’**) Inhibition of Neur by Brd^R^ in the Optix domain (nuclear RFP, red) induced a loss of apical constriction of epi-NSCs (Neur-GFP, green) and persistence of apical E-cad (white) past the epi-NSCs (arrow). (**C**) Analysis of apical area in cells expressing Brd^R^ and in control wild-type cells of the Vsx1 domain, plotted as in Fig 2B showed that inhibition of Neur led to reduced apical constriction (*n* = 7 brains; means ± SEM were calculated using 34-210 cells per binned distance; p = 8×10^−29^, two-tailed student t-test). (**D**,**D’**) Expression of only Neur-resistant isoforms of Sdt (loss of Sdt-GFP3, green), following excision of exon 3 of the *sdt*^*GFP3*^ allele, interfered with apical constriction of epi-NSCs (Neur-GFP, green; E-cad, red). (**E**) Quantification of apical area in cells of the Optix domain expressing Neur-resistant isoforms of Sdt compared to control wild-type cells from the Vsx1 domain, plotted as in Fig 2B, showed the failure to down-regulate Sdt in the epi-NSCs (0-2μm along the x-axis) resulted in intermediate level of apical constriction (*n* = 8 brains, means ± SEM were calculated using 82-319 cells per binned distance; p = 5.9×10^−26^, two-tailed student t-test). (**F-G’’**) While higher levels of junctional MyoII (MyoII-TagRFP, red) were observed at straight interfaces between epi-NSCs (Neur-GFP, green) and more lateral TZ cells in wild-type (F-F’’; arrows), both increased MyoII levels and straight interfaces were less clearly observed when Neur resistant isoforms of Sdt were expressed by *optix-*Gal4 inducing FLP mediated excision of exon 3 of *sdt* (G-G’’). (**H**) Quantification of junctional MyoII along different cell-cell contacts (data plotted as in Fig 2B,C; see Fig 2C for the wild-type control; *n* = 9 brains; ns, non-significant difference, One-way ANOVA).

We next tested whether Neur regulates apical constriction via the regulation of Sdt. To do so, we quantified the apical area of TZ cells expressing only the Neur-resistant isoforms of Sdt, i.e. upon excision of *sdt* exon 3 in the *sdt* allele. An increase in the apical area of epi-NSCs was observed (Fig.5D-E). We therefore conclude that Neur regulates apical constriction in epi-NSCs in part by inducing the Sdt-mediated down-regulation of the Crb complex.

However, the apical constriction phenotype seen upon excision of *sdt* exon 3 appeared to be milder than the one observed upon the inhibition of Neur by Brd^R^ (Fig.5C,E). We therefore suggest that Neur regulates apical constriction via additional targets and/or mechanisms.

Consistent with this interpretation, Neur is known to positively regulate apical constriction and actomyosin contractility independently of Sdt during ventral furrow formation in the embryo (Perez-Mockus et al., 2017a). We conclude that Neur regulates apical constriction in epi-NSCs in part via the down-regulation of Crb.

Finally, we observed that the accumulation of MyoII at the epi-NSC/TZ junction was lost upon excision of exon 3 in the *sdt* allele (Fig.S5). This observation is consistent with the notion that Crb anisotropy promotes the formation of supracellular MyoII cables via the regulated distribution of aPKC (Röper, 2012) and/or Rho kinase (Sidor et al., 2020). Indeed, low aPKC levels were detected at apical junctions of epi-NSCs where junctional MyoII was found to be enriched (Fig.S5) and this distribution of junctional aPKC appeared to depend on the down-regulation of Crb by Neur since expression of the Neur-resistant isoforms of Sdt led to a more uniform distribution of aPKC (Fig.S5). Thus, the down-regulation of Crb by Neur in epi-NSCs not only regulates apical constriction but may also promote the formation of supra-cellular myosin cables via the regulated distribution of aPKC and/or Rho Kinase at epi-NSC junctions.

### RhoGEF3 regulates apical constriction independently of the regulation of Sdt by Neur

A second potential regulator of cortex remodeling in epi-NSCs is RhoGEF3, an exchange factor for Rac/Cdc42 (Couturier et al., 2017; Nakamura et al., 2017). Like Neur, RhoGEF3 is expressed in neural progenitor cells downstream of proneural factors (Couturier et al., 2017), suggesting that RhoGEF3 may also be upregulated in epi-NSCs. Using a knock-in allele of RhoGEF3 endogenously tagged with GFP, we found that RhoGEF3 accumulated at the apical cortex of apically constricted TZ cells (Fig.6A-A’’). To assay the role of RhoGEF3 in the TZ, we studied a *rhoGEF3* knockout allele (Couturier et al., 2017). First, we observed that Patj was similarly down-regulated in wild-type cells and epi-NSCs (Fig.6B,B’). To determine if RhoGEF3 played a role in apical constriction, we measured the apical area of TZ cells and found that loss of *rhoGEF3* led to a partial loss of apical constriction (Fig.6C-D). Thus, RhoGEF3 appeared to contribute to the apical constriction of epi-NSCs independently of the down-regulation of the Crb complex. We also measured apical constriction in *sdt*^*Δexon3*^, *rhoGEF3*^*KO*^ double mutant brains and found that the double mutant had a phenotype similar to each single mutant (Fig.S6). Assuming that Neur-Sdt and RhoGEF3 act within a linear genetically-defined pathway to regulate apical constriction in epi-NSCs, one possible interpretation is that RhoGEF3 acts downstream of Sdt regulation by Neur. Alternatively, RhoGEF3 and Neur-Sdt may act in parallel pathways to regulate apical constriction. Since the loss of *rhoGEF3* had no clear effect on the down-regulation of the Crb complex, we wondered whether MyoII accumulated at the interface between epi-NSCs and other TZ cells in the absence of RhoGEF3. We therefore looked at MyoII-TagRFP and Neur-GFP in cultured *rhoGEF3* mutant brains and found that junctional MyoII remained significantly enriched at the cell-cell contacts between epi-NSCs and their immediate TZ neighbors (b junctions; Fig.6E-F; compare with wild-type brains in Fig.3C). This observation strengthened the view that low Crb, and not apical constriction, directs the formation of supra-cellular actomyosin cables.

**Figure 6:**
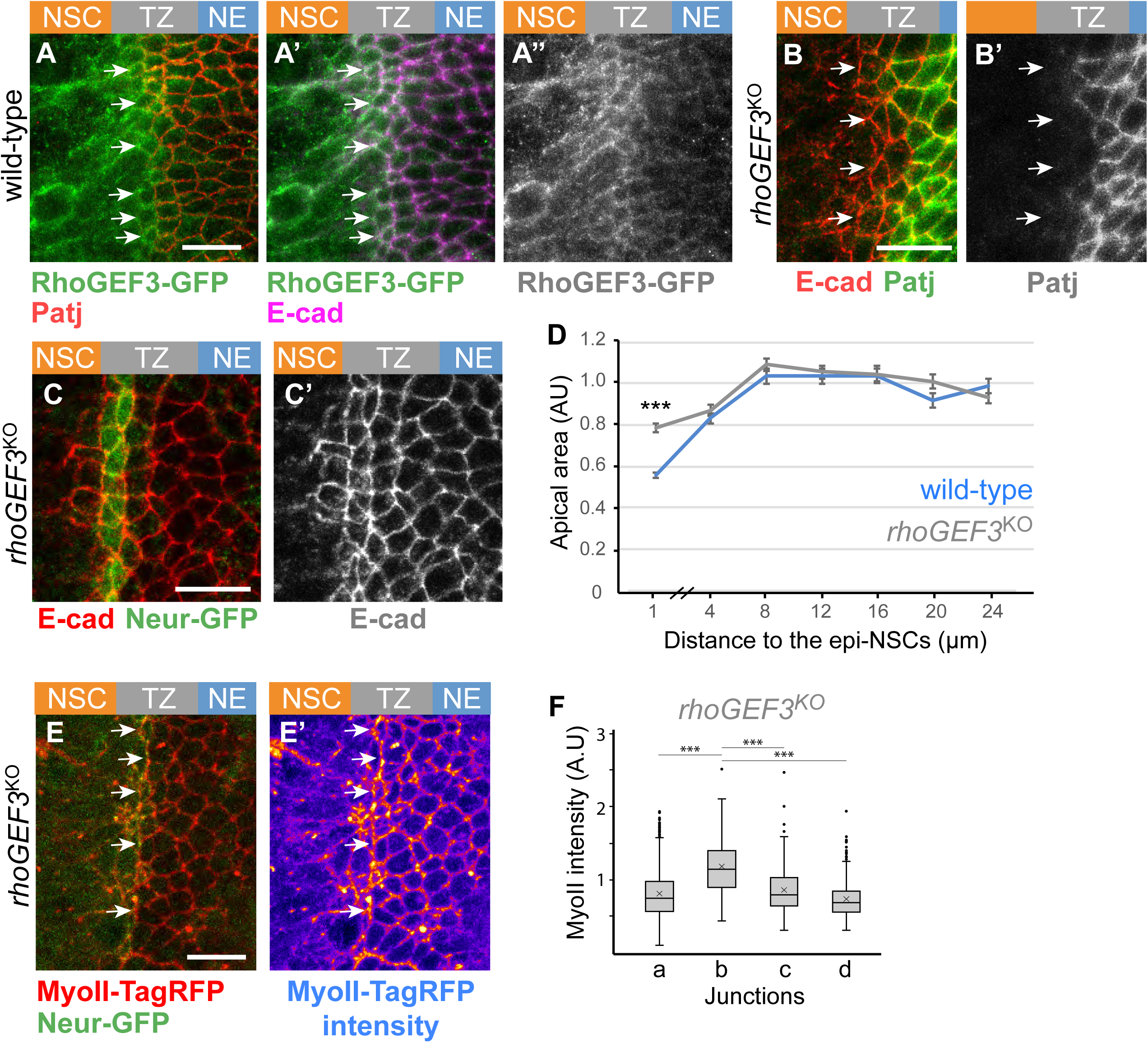
RhoGEF3 is required for apical constriction. (**A-A’’**) RhoGEF3-GFP (green) was detected at low levels at apical junctions of the NE (Patj, red; E-cad, magenta) and appeared to be upregulated in NSCs and epi-NSCs (white arrows, epi-NSCs were identified via the down-regulation of Patj). (**B**,**B’**) Down-regulation of Patj (green) appeared to occur normally in cells located at the medial edge of the NE in *rhoGEF3* mutant brains (white arrows; E-cad, red). (**C**-**D**) Apical constriction of Epi-NSCs (Neur-GFP, green in C) was reduced in *rhoGEF3* mutant brains (C,C’; E-cad, red), as shown in the quantification (D) of the apical area in *rhoGEF3* mutant (*n* = 9 brains) vs wild-type brains (*n* = 7; data presented as in Fig. 2B, with 149-326 cells per binned distance; p = 7×10^−19^, two-tailed student t-test). (**E**,**E’**) Junctional MyoII (MyoII-TagRFP, red) appeared to be enriched along at cell-cell contacts between epi-NSCs (Neur-GFP, green) and more lateral TZ cells (white arrows). (**F**) Box-plots of MyoII intensity along junctions of the NE based on the 4 distinct junction identities described in Fig. 3C, showing that b junctions had a significant enrichment of MyoII in *rhoGEF3*^*KO*^ brains (*n* = 7 brains, b *vs* a, b or c p values <10^−12^, One-way ANOVA, Scheffé multiple comparison tests). Scale bars = 10 μm.

### Coupling fate transition in individual cells with epithelium remodeling at the tissue scale

The dynamics of the NE-NSCs fate transition has so far mostly been considered in one dimension, with the OL epithelium viewed as a cross-section and with a particular focus on temporal dynamics (Egger et al., 2010; Weng et al., 2012; Ngo et al., 2010; Wang et al., 2011; Orihara-Ono et al., 2011). In contrast, how fate dynamics is coordinated in a two-dimensional epithelium has not been examined. In particular, it is not clear whether all TZ cells juxtaposed to the epi-NSCs progress synchronously to form a new row of epi-NSCs, hence replacing the pre-existing epi-NSCs in a quasi-simultaneous manner, leading to a saltatory progression of the epi-NSC front, or whether individual TZ cells become epi-NSCs in a more continuous manner, such that the epi-NSC front progressed in a gradual manner.

To distinguish between these two possibilities, we followed the dynamics of TZ cell entry into the row of epi-NSCs live in cultured L3 brains. Neur-GFP and E-cad-mCherry were used to mark epi-NSCs and to outline epithelial cells, respectively. Tracking all epithelial cells adjacent to the row of epi-NSCs, we monitored these TZ cells as they become epi-NSCs, i.e. express Neur-GFP (Fig.7A-D; a threshold value of Neur-GFP intensity was used to define cells as epi-NSCs). We found that the number of TZ cells becoming epi-NSCs increased at a constant rate over time (Fig.7E). This showed that rows of TZ cells do not adopt synchronously an epi-NSC state in a periodic manner but rather that individual TZ cells switch fate and integrate the row epi-NSCs one cell at a time in a continuous manner.

**Figure 7:**
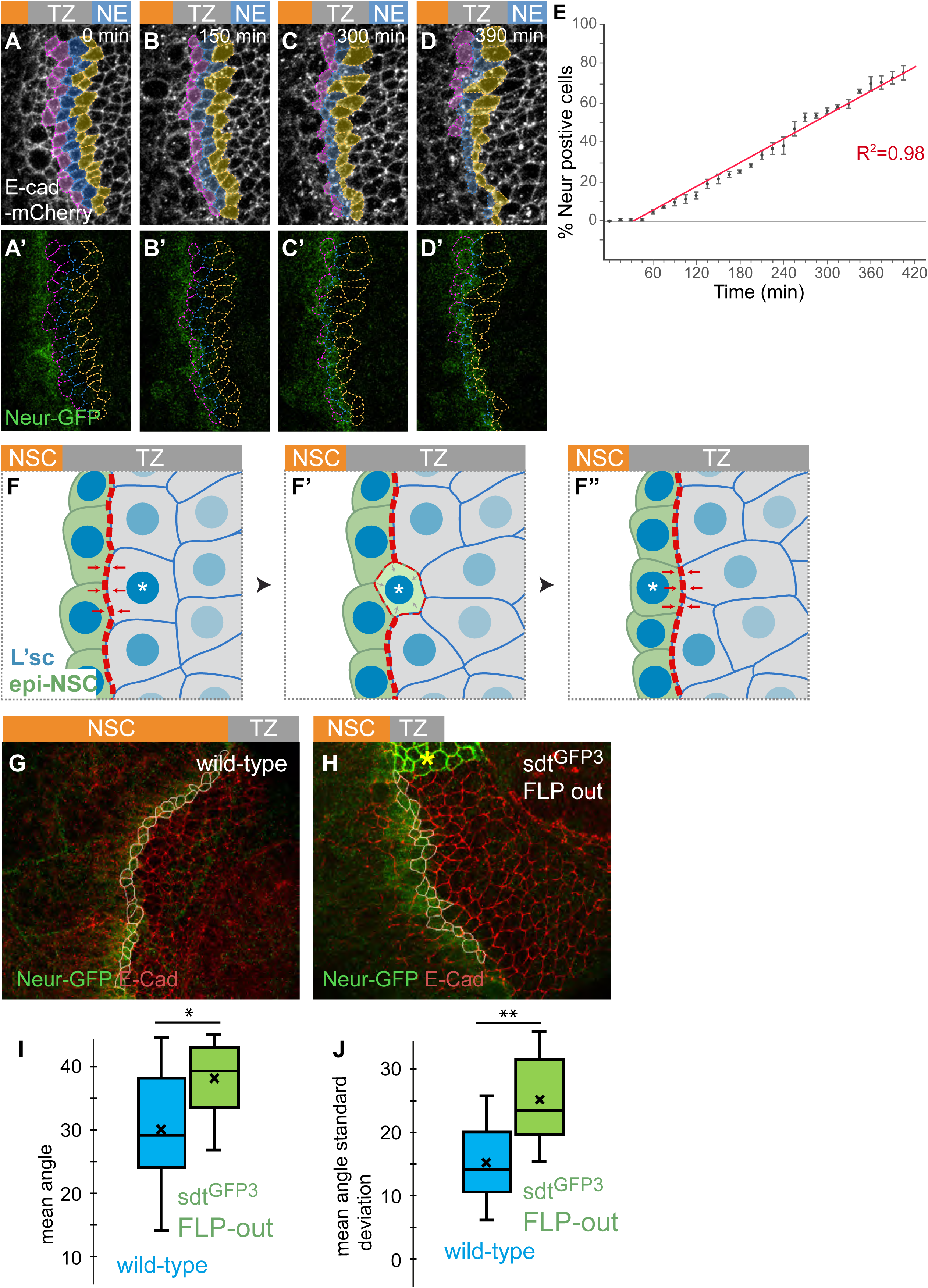
Dynamics of of TZ-epi-NSC transition in space and time. (**A-D’**) Snapshots from a time-lapse movie showing the dynamics of TZ-epi-NSC transition (E-cad-mCherry, white; Neur-GFP, green). TZ cells that have not yet become epi-NSCs were manually tracked (color code refers to the distance to epi-NSCs) (**E**) Cumulative plot of TZ cells switching to the epi-NSC state, based on Neur-GFP expression over time (switch was defined by a threshold of 1 standard deviation below the mean of Neur-GFP intensity in apically constricted epi-NSCs; *n* = 3 brains; 34-47 cells tracked per brain). Transition appeared to be a continuous process. (**F-F’’**) Model of mechanical coupling by Neur. Acquisition of the epi-NSC state (green; L’sc levels in blue) is a probabilistic event occurring at the single cell level (F). It is associated with the expression of Neur that in turn induces apical constriction (F’). Neur down-regulates Crb in a cell-autonomous manner, hence moving the boundary between low and high Crb cells, hence the enrichment of junctional MyoII (red dotted line) resulting from the crb anisotropy: MyoII accumulation increases at the lateral junction and decreases at the medial junction (arrows in F’). This is proposed to modify tension along this interface and promote cell-cell rearrangement to direct integration of the newly-fated epi-NScs into the one-cell row of epi-NSC at the medial edge of the TZ (F’’). This mechanical coupling may ensure precise tissue architecture and regular spatial progression of the differentiation front. (**G**,**H**) Immunostainings showing the epi-NSCs (Neur-GFP, green; E-cad, red; cell outlines in white) located at the interface with the Neur-negative TZ cells which were used to evaluate precision of morphogenesis (G: wild-type; H: *optix-Gal4 UAS-flp sdt*^*GFP3*^: Neur-resistant cells do not express Sdt-GFP3, green; yellow star) (**I**,**J**) Plots showing the mean angle values (I; n=12 wild-type brains and n=10 *optix-Gal4 UAS-flp sdt*^*GFP3*^ brains; p=0.019, two-tailed Student t-test) and standard deviations (J; p value =0.002, two-tailed Student t-test) showing that medial epi-NSCs formed a less regular row of cells upon excision of exon 3 of sdt, indicating that regulation of Sdt by Neur contributes to the integration of Neur-positive TZ cells into the row of epi-NSCs.

Our observations therefore suggest a model whereby fate transition is mechanically coupled with cellular rearrangement to promote a smooth progression of the differentiation front (Fig.7F-F’’). Specifically, we propose that the up-regulation of Neur in induces apical constriction and down-regulation of Crb via Sdt (Fig.7F’). As a consequence, a new low-high Crb interface is created laterally, with the old low-high Crb cell-cell contact being converted into a low-low Crb interface. These changes would then impact on the junctional accumulation of MyoII and reposition the cell-cell contacts that are under tension, thereby promoting the integration of the newly-specified epi-NSC into the row of pre-existing epi-NSCs (Fig.7F’’). This model predicts that a failure to down-regulate Crb in cells progressing towards the epi-NSC state should result in defects in the integration of these cells into the row of epi-NSCs. To test this prediction, we evaluated this integration in wild-type brains (Fig.7G) and examined the effect of uncoupling fate specification and Neur expression from Crb down-regulation upon excision of the exon 3 of the *sdt*^*GFP3*^ allele (Fig.7H). To do so, we measured the angles between the lines linking the centroid of each medial epi-NSCs to those of its two neighbours relative to the local NE-to-NSC front. We found that the loss of Crb down-regulation by Neur in cells expressing Neur-resistant Sdt isoforms led to higher mean angle values (mean per brain) compared to wild-type controls (Fig.7I). It also increased cell-to-cell variability within each brain as seen by the difference in mean standard deviation values (mean per brain; Fig.7J). These data indicate that the regulation of Sdt by Neur contributes to the proper integration of the TZ cells becoming epi-NSC into the epi-NSC domain. We therefore suggest that the loss of Crb by Neur regulates cell-cell rearrangement via its effect on junctional MyoII to facilitate the integration of the TZ cells progressing towards the epi-NSC state into the epi-NSC domain. Thus, adoption of the NSC fate appears to be intrinsically coupled with the mechanical changes at apical junctions that are induced by the Neur-dependent down-regulation of Crb, hence promoting the formation of a continuous single-cell row of epi-NSCs.

## Discussion

The emergence of NSCs from the NE of the OL provides a good model to study how acquisition of a stem cell identity is associated with changes in epithelial tissue organization. Here we identify a transient cellular state during the NE-to-NSC transition that we named the epi-NSC state. It is characterized by the persistence of epithelial features, including apical E-cad, apical constriction, expression of Neur and loss of Crb. We find that TZ cells are in the G2 phase of the cell cycle and that epi-NSCs enter mitosis and divide asymmetrically to produce the first-born medulla neurons of the NSC lineage. As individual TZ cells acquire this epi-NSC state and apically constrict, each emerging epi-NSC individually integrates the epi-NSC row. The cell density along the differentiation front is higher than those of NE but no invagination forms, indicative of cell-cell intercalation. Neur in epi-NSCs regulates not only Delta signaling but also apical constriction, in part via the down-regulation of Crb. Low Crb in epi-NSCs creates an anisotropy in Crb levels within the TZ. This anisotropy appears to result in the formation of supra-cellular MyoII cables. Down-regulation of Crb by Neur is required for the proper integration of these emerging epi-NSCs into the row of previously-established epi-NSCs. Thus, mechanical coupling by Neur is proposed to link fate specification with the formation of a continuous stripe of precisely lined-up epi-NSCs in the OL.

Recent studies have highlighted that Neur not only regulates Notch signaling via the endocytosis of Delta but also contribute to the regulation of epithelium morphogenesis via its interaction with Sdt. These two activities of Neur coincide in the TZ of the OL to coordinate fate and cell shape changes during the NE-to-NSC transition. Indeed, Neur is not only required for Notch activity at the TZ (Contreras et al., 2018) but also down-regulates Crb in epi-NSCs via specific Sdt isoforms. Analysis of cells expressing only Neur-resistant Sdt isofomrs showed that this activity of Neur is important for apical constriction. Neur was also proposed to regulate via an unknown molecular target the contractile activity of actomyosin at gastrulation. Whether Neur similarly regulates MyoII dynamics in epi-NSCs to promote apical constriction remains to be studied. However, since inhibition of Neur has a stronger effect on apical constriction than blocking the regulation of Crb by Neur, we suggest that Neur regulates apical constriction only in part via Sdt. In the ratchet model of apical constriction, first proposed for the apically constricting cells of the ventral furrow in the early fly embryo (Martin et al., 2009), apical constriction is induced by the contractions of the medial actomyosin meshwork pulling onto apical junctions, combined with a rapid remodeling of the apical junctions dissipating the energy stored (Clement et al., 2017; Mason et al., 2013; Martin et al., 2010). Since MyoII pulses were observed in both NE and epi-NSCs, our observation that only epi-NSCs, not NE cells, undergo apical constriction in response to repeated pulses of medial MyoII may suggest that the apical cortex is remodeled faster in epi-NSCs than in NE cells, hence contributing to stabilize a loss of apical area upon constriction (Clement et al., 2017). For instance, E-cad could be internalized at a faster rate in the absence of Crb in epi-NSCs. While it is not known whether the kinetics of E-cad endocytosis vary between NE and epi-NSCs, we note that this view is entirely consistent with the notion that Crb stabilizes AJs and is required to maintain epithelium integrity in contexts of morphogenesis, e.g. cell delamination, cell-cell intercalation and division (Harris and Tepass, 2008; Bajur et al., 2019; Flores-Benitez and Knust, 2015; Campbell et al., 2009; Perez-Mockus et al., 2017b; Röper, 2012; Salis et al., 2017). Future studies will address whether Neur promotes a faster turn-over of E-cad via its regulatory activity on the Crb complex and/or cortical actomyosin.

Whether Neur similarly regulates the delamination of NSCs from the neuroectoderm in the early fly embryo is an interesting possibility that remains to be studied (Perez-Mockus and Schweisguth, 2017). In the embryonic neuroectoderm, delaminating neuroblasts exhibit down-regulation of both Crb and E-cad preceding the first round of asymmetric division (Simões et al., 2017). In contrast, Crb but not E-cad are down-regulated in epi-NSCs in the OL prior to division. Whether this difference in E-cad distribution merely reflects a difference in geometry, e.g. delamination of individual cells surrounded by ectodermal cells which reinforce their AJs in response to Notch versus progressive and collective changes along a front, or molecular differences in regulatory mechanisms remain to be explored.

Interestingly, while Neur promotes EMT, Notch has been shown to prevent mesenchymal transition in fly imaginal disc epithelia (Boukhatmi et al., 2020) and human lung cancer (Ito et al., 2017). This raises the possibility that high Notch activity in the TZ blocks the NE-to-NSC transition in all TZ cells prior to the Notch ON/OFF switch. Thus, Neur in epi-NSCs could not only regulate an EMT-like process in epi-NSCs but also contribute to inhibit this same process in its TZ neighbors by promoting high Notch activity. By extension, NSC delamination in the embryonic neuroectoderm can be seen as a collective process involving both the delaminating cell and its ectodermal neighbors as delaminating NSCs signal to their neuroectodermal neighbors to remain epithelial. Whether and how Notch activation in neuroectodermal cells regulate in a feed-back manner NSC delamination remains to be studied (An et al., 2017; Arefin et al., 2019).

Neur is probably only one of several factors expressed in TZ cells downstream of the proneural activator L’sc and involved in the regulation of apical constriction and cell-cell rearrangements. Here, we identify one such factor, the Cdc42/RacGEF known as RhoGEF3 (Couturier et al., 2017), that may act in parallel to Neur. Thus, while genetic analysis strongly suggests that the regulation of Crb by Neur is not essential for proper development, Neur may either act redundantly with other regulatory mechanisms of epithelium morphogenesis and/or ensure efficient and coordinated tissue development. In the OL, we find that regulation of Crb by Neur contributes to line-up epi-NSCs. Mechanistically, this Neur-dependent cell-cell rearrangement appears to be mediated by a change in the distribution of junctional MyoII as Crb levels go down.

Reproducibility in morphogenesis is commonly observed and often involves at least two mechanisms. One relies the precise decoding of positional information by gene regulatory networks (Petkova et al., 2019), with output genetic programs regulating cell shape changes at the single cell level. In the context of the NE-to-NSC transition, NE cells presumably read their distance to the NSC front through a combination of EGFR and Notch signalling to regulate L’sc, but how EGFR and Notch contribute to the precise temporal and spatial dynamics of L’sc remains to be studied. The other involves tissue-scale mechanical coupling to organize cells in space in a reproducible manner (Eritano et al., 2020; Aliee et al., 2012; Yevick et al., 2019). For example, neural progenitors in the fish neural tube rearrange cell-cell contacts to establish sharp boundary through cell sorting. This sorting process corrects imprecision of spatial patterning through spatial averaging and contributes to establish shar boundaries (Xiong et al., 2013). Our analysis of the Neur-dependent regulation of Crb in epi-NSC suggests that Crb anisotropy within the TZ results in the formation of supracellular MyoII cables that contributes to precisely line-up cells in a similar epi-NSC state along a continuous row. Accordingly, mechanical coupling of the Neur-expressing cells with low Crb contributes to define a spatially precise NSC front. Indeed, any stochastic variations in the kinetics of cell fate transition, as TZ cells dynamically progress towards the epi-NSC state, is a potential source of imprecision. However, correlated variations in the kinetics of increased Neur levels are proposed to be spatially buffered by mechanical coupling: precocious high Neur is translated into a spatial shift in the boundary of Crb anisotropy, leading to a change in junctional MyoII accumulation and cell-cell rearrangement, so that any TZ cell with higher Neur compared to its immediate neighbor is integrated first into the row of epi-NSCs. Thus, regulation of Crb by Neur in the TZ contributes to coordinate fate acquisition at the single cell level with epithelium remodeling at the tissue level to ensure a smooth progression of the differentiation front. While it is not known whether precision of this particular morphogenetic process is relevant for fitness, we speculate that this may help in the precise mapping to photoreceptor cells being simultaneously specified in the developing retina (Sato et al., 2013). Future studies will address whether Neur operates in other developmental contexts to regulate both Notch-dependent fate dynamics with changes in cell polarity and cortex contractility to mechanically couple at the tissue scale cells that are in a similar state and thereby ensure precision in morphogenesis.

## Acknowledgements

We thank Y. Bellaïche, A. Brand, Y. Cai, L. Cheng, D. Glover, Y. Hong, Y. N. Jan, M. Krahn, F. Matsuzaki, A. Salzberg, M. Sato, J. Skeath, M. Suzanne, A. Wodarz, the Developmental Studies Hybridoma Bank (DSHB), Flybase and Image Analysis Hub (IAH) of the Institut Pasteur for reagents and resources. We thank S. Herbert (IAH), R. Levayer, J.-Y. Tinevez (IAH), T. Schweisguth and L. Valon for help in segmentation, local z-projection and image data analysis, M. Rujano for help in live imaging and L. Couturier for technical help. We thank S. Chanet, E. Contreras, A. Hakes and G. Perez-Mockus for critical reading. This work was funded by the ANR-10-LABX-0073 and FRM-DEQ20180339219 grants.

## Materials and methods

### Flies

*D. melanogaster* flies were raised in food vials at 25°C and larvae were allowed to develop until the desired stage. For all RNAi experiments embryos and larvae were raised at 29°C instead of 25°C. Wild-type controls were *w*^*1118*^ flies. Otherwise, the following alleles and transgenes were used: *E-cad-mcherry* (Herszterg et al., 2013) (from Y. Bellaiche); *crb-GFP-A* (Huang et al., 2009)(from Y. Hong); *sdt*^*GFP*^, tagging all Sdt isoforms (Perez-Mockus et al., 2017b)); *sdt*^*GFP3*^, a GFP tagged and FRT flanked knock-in allele of the *sdt* gene tagging all long isoforms of Sdt containing the NBM (Perez-Mockus et al., 2017b)) ; *sdt*^*Δ3GFP*^, a mutant allele with a deletion of the GFP-tagged exon 3 from the *sdt*^*GFP3*^ allele (Perez-Mockus et al., 2017b) ; *rhoGEF3*^*KO*^ (Couturier et al., 2017); MyoII-GFP and MyoII-TagRFP CRISPR knock in alleles (Gracia et al., 2019) (from M. Suzanne) also noted *sqh*^*eGFP*^ and *sqh*^*Tag-RFP*^; Neur-GFP (Perez-Mockus et al., 2017a) ; RhoGEF3-GFP (Couturier et al., 2017) ; aPKC-GFP (Besson et al., 2015) ; ubi-Par3-mCherry (Herszterg et al., 2013) (from Y. Bellaïche; also noted Baz-Cherry) ; *optix-*Gal4 (Gold and Brand, 2014)(NP2631-Gal4; from A. Brand); *hh-*Gal4; FRT82B *neur*^*IF65*^; UAS-Brd^R^ (a transgene in which K residues were mutated into R residues was used to drive ectopic expression of a stabilized version of the protein Brd (Perez-Mockus et al., 2017b)) ; UAS-mib1 (Le Borgne et al., 2005) ; ubi-GFP-E2F1 combined with ubi-mRFP-CycB (FUCCI, BL-55124) ; UAS-dsRNAi *l’sc* (BL-51443) ; Histone2Av-RFP (His2Av-RFP; BL-23651) and UAS-flp. The following stocks were used for MARCM clone experiments: *hs-flp UAS-nlsGFP tub-Gal4*;; FRT82B *tub-Gal80*/TM6B and *hs-flp UAS-myrRFP tub-Gal4*;; FRT82B *tub-Gal80*/TM6B. Larvae were heat-shocked for 1 hour at 36.5°C to induce *flp* expression.

### Immunostaining

Third larval brains were dissected in PBS and fixed in 4% formaldehyde/PBS on shaker for 25 min. The brains were then washed 3 × 5 min with 0.3% Triton X-100 in PBS (PBST), and incubated on a shaker overnight at 4°C with primary antibodies. The brains were then washed 2 × 10 min with PBST and incubated with secondary antibodies for 3h at room temperature or overnight at 4°C. After secondary antibody incubation and washes, brains were mounted in 4% *N*-propylgalate and 80% glycerol. The following antibodies were used: rabbit anti-aPKC (1:500; Santa Cruz Biotechnology, Inc.), guinea pig anti-Patj (1:500; a gift from M. Krahn), rabbit anti-Par3 (1:500; a gift from A. Wodarz), rat anti-DCAD2 E-Cad (1:200; from the DSHB), guinea pig anti-Dpn (1:5,000; a gift from J. Skeath), guinea pig anti-L’sc (1:1,200; a gift from M. Sato), mouse anti-Wor (1:1,000; a gift from Y. Cai), rabbit anti-Ase (1:5,000; a gift from Y. N. Jan), rabbit anti-Hth (1:1,000 ; a gift from A. Salzberg), mouse anti-Pros (1:50 ; MR1A from the DHSB), rabbit anti-Mira (1:1,000; a gift from F. Matsuzaki), mouse anti-pH3 (1:400; Cell Signaling, Inc.), rabbit anti-CycB (1:500; a gift from D. Glover), rabbit anti-DsRed (1:200, Clonetech), goat anti-GFP (1:500; Abcam). All secondary antibodies were Alexa Fluor 488–, Cy3-, and Cy5-coupled antibodies from Jackson ImmunoResearch Laboratories, Inc and were diluted 1:500 in PBST. Brains were stained for actin using atto647N-phalloidin (1:1,000; Sigma-Aldrich). Brains were stained for DNA using DAPI (1μg/ml, Sigma).

### Confocal imaging and image data analysis

Images were acquired using a confocal microscope (LSM780; ZEISS) with a 63× Plan Apochromat 1.4 NA differential interference contrast M27 objective. Cross-section views actually correspond to a single plane from a confocal z-stack taken as a lateral image of the NE. Images showing the nuclear layer of the NE were maximal projections of two to five planes from confocal z-stacks (Δz = 1 µm) using Fiji (Schindelin et al., 2012). Surface views of the NE were either max projections (3 planes from a z-stack, Δz = 1 µm) or local z projections performed using Fiji. This facilitated constructing a flattened view of the curved epithelia for analysis. For local z projections the following threshold and post filtering were specified: Neighbourhood size = 30 pixels, binning = 1, Gaussian filter = 0, median post-filter size = 15 (to facilitate curvature smoothening) and Δ*z* = 2-3 were used. The E-cad signal was used as the reference plane for local z projections. Distances from the TZ were calculated by segmenting cells based on apical membrane signals (E-cad). E-cad signal was first smoothed by using the ‘remove outliers’ tool and then by the Median filtering 3D tool in Fiji. The E-cad signal was then projected using the smooth manifold extraction plugin in Fiji. Cells were then segmented (Watershed segmentation) using the TissueAnalyzer plugin for Fiji and hand corrected. Cell distances to the TZ were calculated in MATLAB by measuring the shortest distance of cell centroids to ROIs drawn along apically constricted epi-NSCs expressing Neur. To estimate integration of Neur-positive TZ cells into the row of epi-NSC, we first segmented epi-NSCs based on E-cad and Neur expression in TissueAnalyzer from max-projected images and identified cell centroid coordinates in the tissue. We then measured and summed for each medial epi-NSCs the values of the two angles formed by the local NSC front (local linear regression of 5 centroid positions) and a line linking its centroid and to those of its immediate epi-NSC neighbour, left and right (angles for 10-35 epi-NSCs per brain were measured; mean and standard deviation values were then plotted in Figures 7I,J).

### *ex vivo* brain culture

Third instar larval brains were dissected in Schneider’s insect media (Sigma-Aldrich) with 5% Fetal Bovine Solution (FBS, Gibco). Larval brains were then mounted in Fibrinogen (MP Biomedicals; 5 µg diluted in 1mL of Schneider’s insect media solution) and clotting was induced using thrombin (MP Biomedicals) in 35 mm MatTek glass bottom dishes. Brains were then cultured in 2 mL of Schneider’s insect media containing 5% FBS, 0.5% penicillin (Gibco), Ecdysone (20 nM) (Sigma-Aldrich) and insulin (200 µM) (Sigma-Aldrich). Cultured brains were live imaged for a period between 1-7 hours on a confocal LSM780 (Zeiss).

### Live imaging and image processing

To measure apical constriction and MyoII pulses, the following conditions were used: (Δ*t* = 15s intervals, Δ*z* = 1 μm, 20 *z*-sections) movies acquired with a 63× Plan Apochromat 1.4 NA differential interference contrast M27 objective. For segmentation, the movies were first locally z-projected using the E-cad-mCherry or Par3-mCherry signal as the reference plane. The following threshold and post filtering were specified: Neighbourhood size of 21 pixels, binning of 4, median post-filter size of 41 and Δ*z* = 2 were max projected. Cells located at the medial edge of the NE that were observed to apically constrict during the course of the movie as well as more lateral cells of the NE were hand-segmented at each time point using the TissueAnalyzer plugin for Fiji. We used custom-made programs for quantification and analysis. For each cell, apical area measurements were smoothed using a Gaussian filter (sigma value was 1.5x time step, i.e. ∼22 sec). These values were then used to derive apical area rate of change: Δarea = area (t^n+1^) – area (t^n^). A contraction phase was defined as Δarea < 0. To identify contraction rate maxima a threshold was set to exclude maxima that fell below the mean constriction rate for that cell in order to filter out noise.

To measure the medial–apical pool of MyoII, the segmentation mask was 1-pixel wide (1 pixel = 1 μm) and only signal that fell within the segmentation mask was included. MyoII intensity was quantified as the mean pixel intensity per cell per time point. For each cell MyoII intensities were smoothed using a Gaussian filter (sigma value was 1.5x time step, i.e.

∼22 sec). These values were then used to derive medial myosin intensity rate of change: ΔMyoII intensity = MyoII intensity (t^n+1^) – MyoII intensity (t^n^). To identify peaks that would distinguish a phase of maximal accumulation of MyoII a threshold was set such that peaks with amplitude values inferior to 25% of the mean of all peaks for each cell were discarded to filter out background noise.

Junctional MyoII was measured from snapshots of *ex vivo* cultured brains. The junctional MyoII signal was quantified in Fiji from max projections. Neur GFP was used to identify epi-NSCs and junctional MyoII (MyoII-TagRFP) was measured by hand tracing ROIs 2 pixels wide along junctions (1 pixel = 1µm) and quantifying mean intensity.

Cross-correlation coefficients (R) between Δarea and ΔMyoII were determined by first identifying time points associated with maximal rates of constriction (thresholded as described above). We then considered an interval of 10 time points (∼150 sec) and plotted the correlation coefficients between the two variables at each time point (for the plots, t0 sec offset = time point of maximal peak of ΔMyoII). The correlation plots for each contraction phase were then merged to create an average cross correlation plot per cell (between 8 to 40 contraction phases were identified per cell).

The average offset time between a max ΔMyoII and a max Δarea contraction were determined by first identifying the time points associated with maximal rates of area contraction (thresholded as described above). For each of these time points the closest associated max ΔMyoII in time was identified (thresholded as described above) and the time offsets between the two maxima were calculated per contraction phase per cell. These offsets were then averaged per cell and pooled for all epi-NSCs and NE cells respectively.

To monitor the dynamics of epi-NSC state acquisition, cultured brains were imaged using: Δ*t* = 15min intervals, Δ*z* = 1 μm and 20 *z*-sections. Images were first locally z-projected using the E-cad signal as the reference plane. The following threshold and post filtering were specified: Neighbourhood size = 30 pixels, binning = 3, Gaussian filter = 0, median post-filter size = 15 (to facilitate curvature smoothening) and Δ*z* = 2 were used to locally max project.

Neur-GFP and E-cad-mcherry were used to define epi-NSCs and NE membranes respectively. Approximately 3 rows of NE cells adjacent to the Neur positive epi-NSCs at T0 were tracked over the time lapse. At each time point cells were segmented manually in Fiji and the average Neur intensity was measured. Cells were defined as having acquired an epi-NSC state when their average Neur intensity signal surpassed a threshold of 1 SD below the mean Neur intensity of all apically constricted epi-NSCs at the medial edge.

The junctional MyoII signal was quantified in Fiji from local z projections by hand tracing ROIs 2 pixels wide along junctions (1 pixel = 1µm).

### Statistical analysis

For statistical calculations R Studio was used to perform Student’s t-tests (two tailed) to compare the means between two samples and one-way ANOVAs to compare the means of three or more samples, where Scheffé multiple comparison tests were then used to perform post-hoc analysis between the means.

## Supplementary Figure legends

**Supplementary Figure 1:**
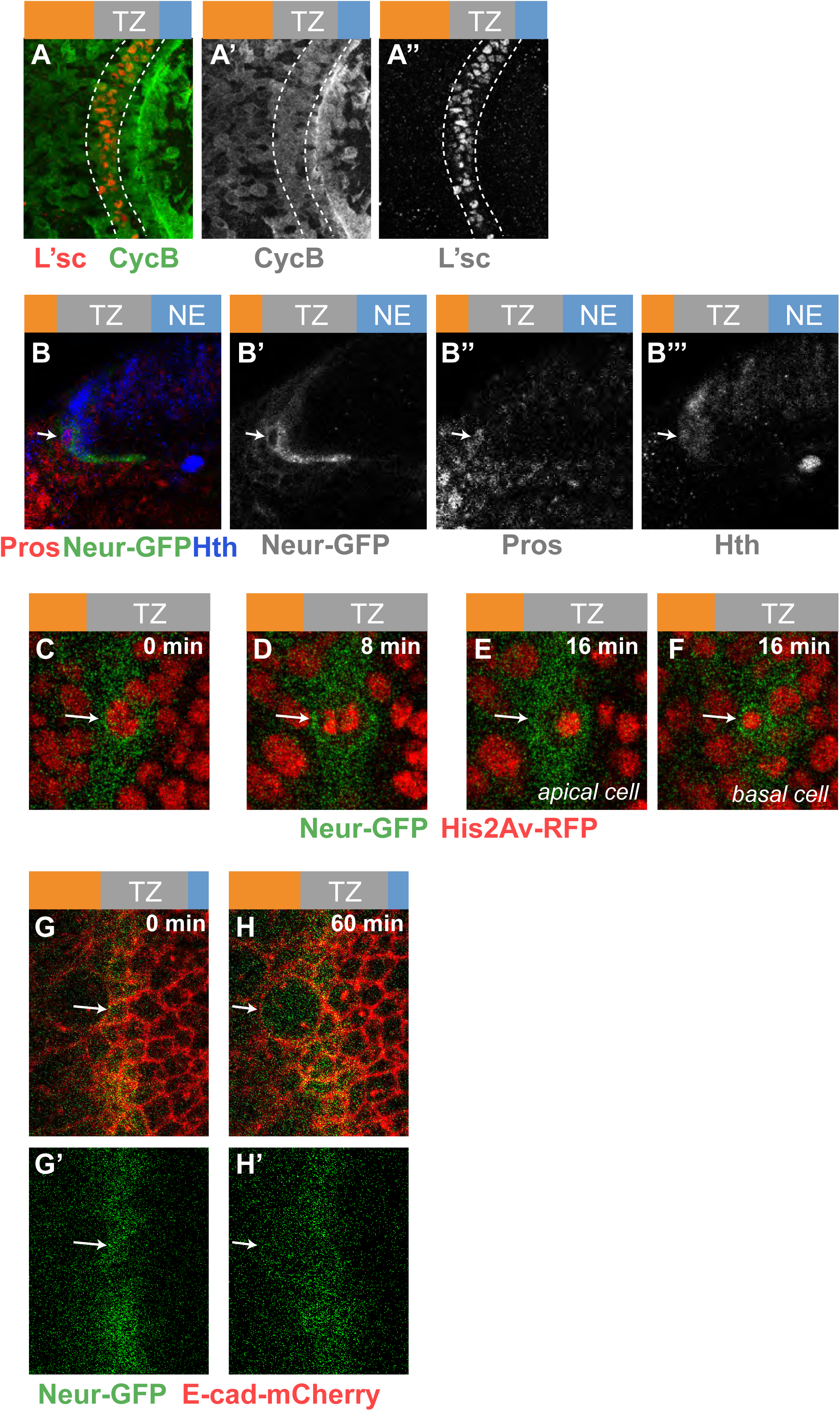
(**A-A’’**) TZ cells (L’sc, red) expressed CycB (green), supporting that they are in G2 phase. In contrast, heterogenous CycB expression is seen in the NE. (**B-B’’’**) Cross-section view showing a small basal cell at the medial edge of the TZ (arrow) expressing Pros (red), Neur-GFP (green) and Hth (blue). This cell likely corresponds to a GMC derived from the asymmetric division of an epi-NSC. (**C-F**) Snapshots of a time-lapse video tracking an epi-NSC (Neur-GFP, green) entering mitosis (His2Av-RFP, red) and dividing asymmetrically to producing a large apical cell (E) and a small basal cell (F). Arrows point to the dividing epi-NSC and its progeny cells. (**G-H’**) Snapshots of a time-lapse video showing a surface view of the TZ (E-cad-mCherry, red). E-cad is retained apically as the epi-NSC (Neur-GFP, green; arrow) undergoes mitotic rounding.

**Supplementary Figure 2:**
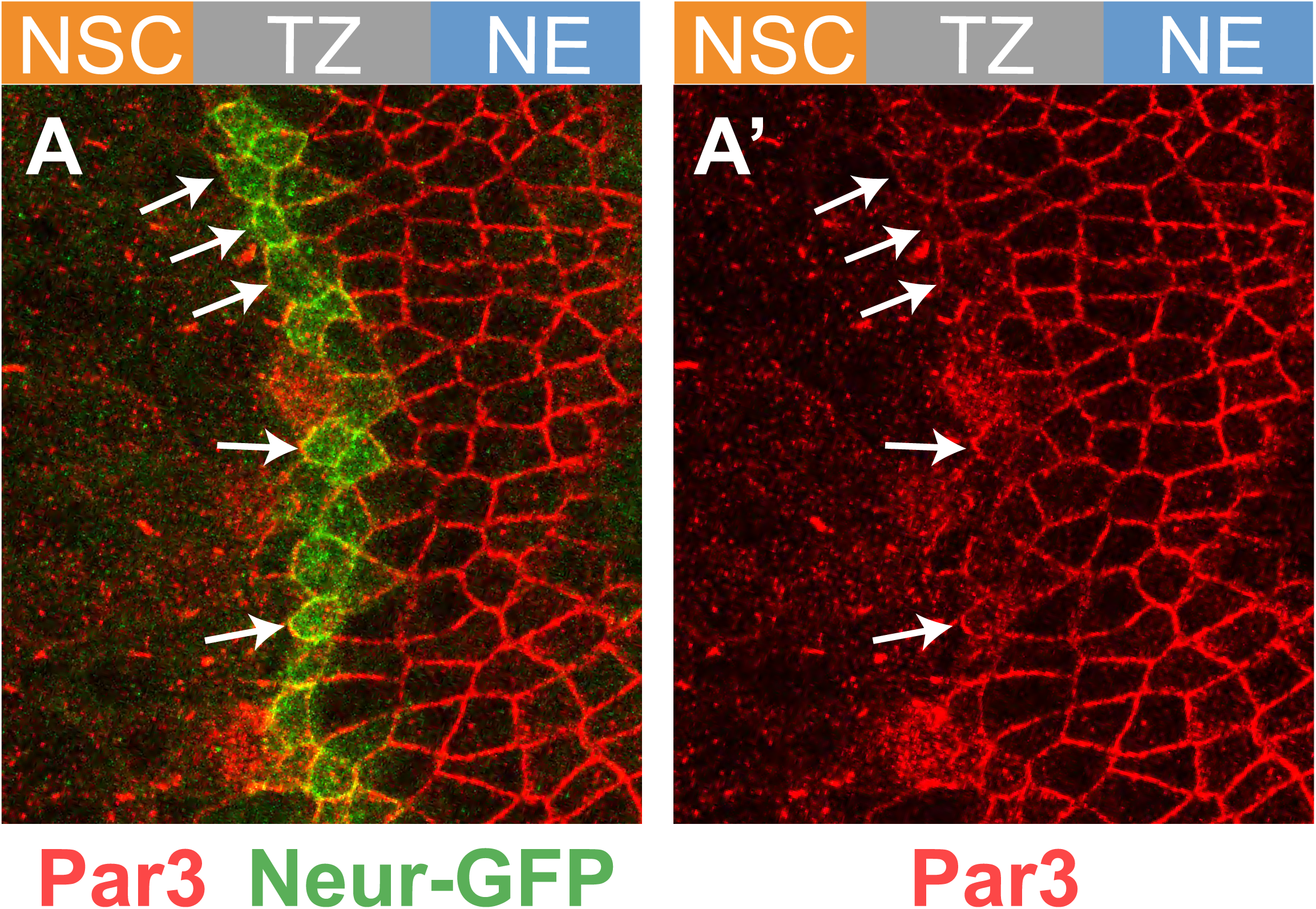
(**A**,**A’**) epi-NSCs (Neur-GFP, green) retain Par3 (red) at junctions of (arrows), albeit at a slightly lower level consistent with junction remodeling in epi-NSCs.

**Supplementary Figure 3:**
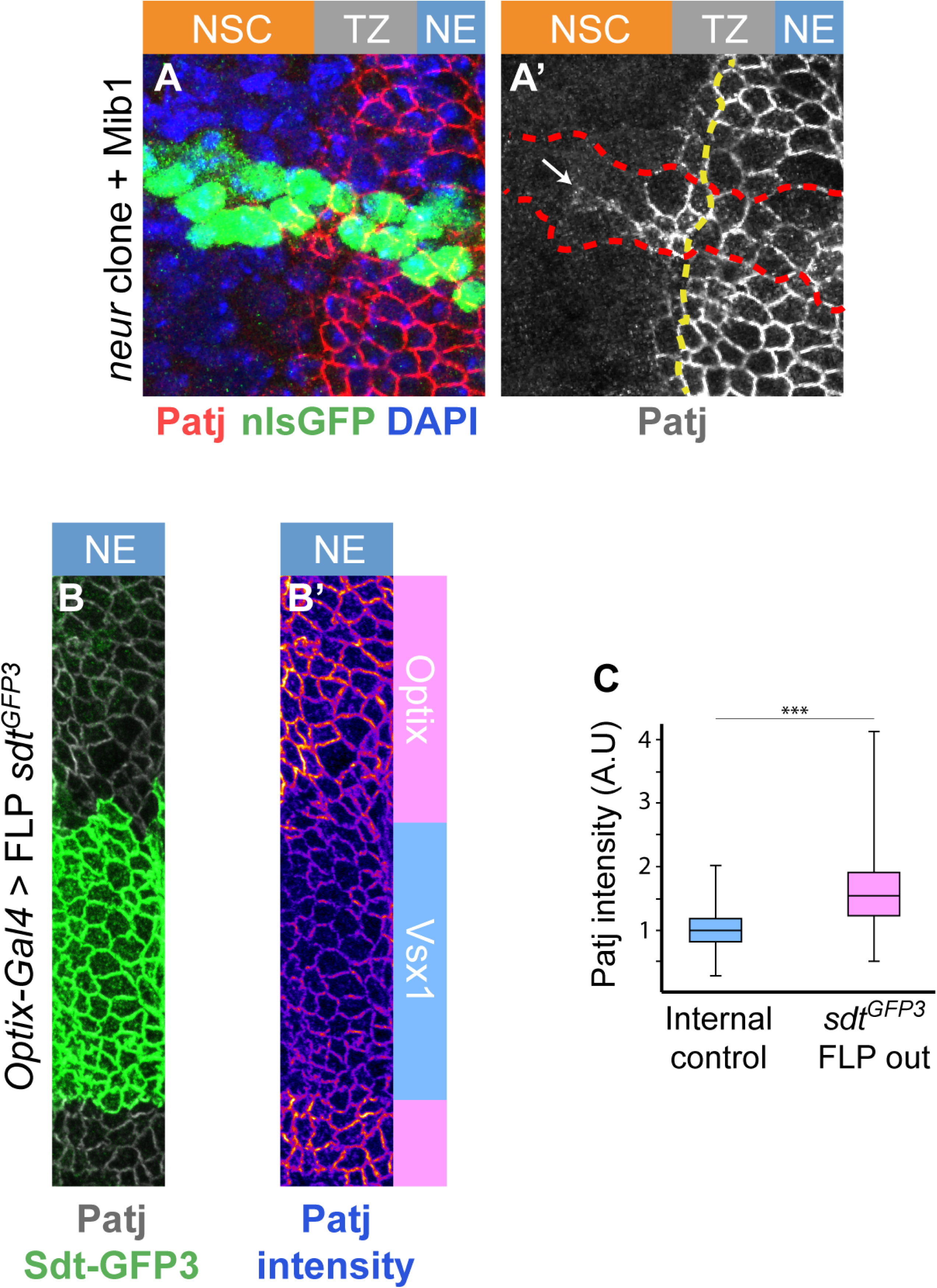
(**A**,**A’**) Patj (red) was observed to persist at the apical cortex of *neur*^IF65^ mutant cells over-expressing Mib1 using *tub-Gal4* past the TZ medial edge (MARCM clones marked by nuclear GFP, green, outlined in red; DAPI, blue). The boundary between epi-NSCs and other TZ cells is shown as a yellow dotted line. (**B-C**) Loss of the long Sdt isoforms in the Optix domains (loss of Sdt-GFP3, green) resulted in slightly higher accumulation of Patj (white, B; intensity code, B’), as quantified in C (n=4).

**Supplementary Figure 4:**
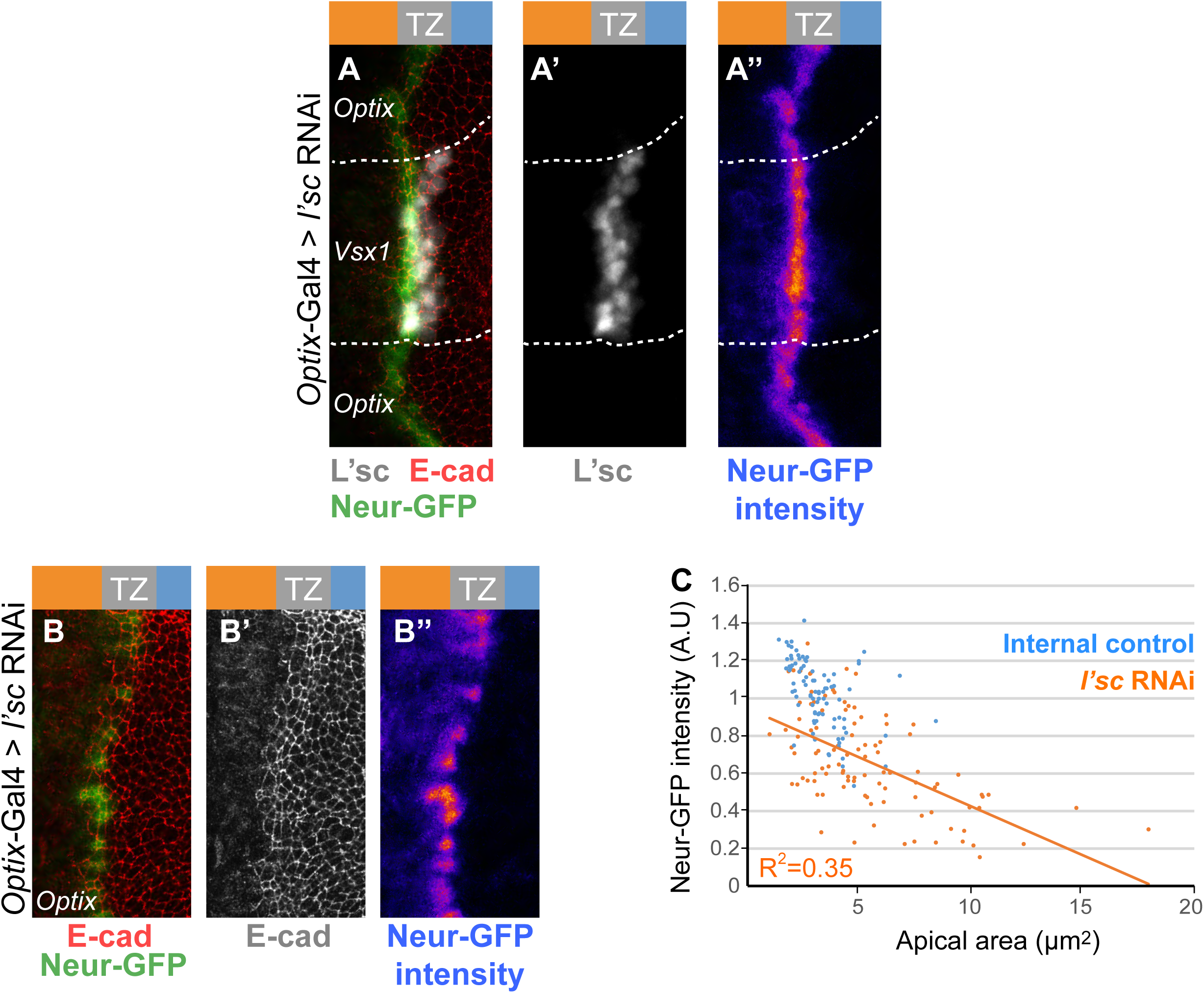
(**A-B’’**) RNAi knockdown of *l’sc* expression via *optix*-Gal4 induced a strong loss of L’sc (A-A’’; white) which was associated with a decrease in Neur-GFP (green in A,B; compare epi-NSCs of the Optix and Vsx1 domains in A-A’’), confirming that L’sc regulates the expression of the *neur* gene. A high degree of cell-to-cell heterogeneity in Neur-GFP levels was seen upon l’sc knock-down in the Optix domain (B’’). (**C**) Neur-GFP intensity values (normalized to mean GFP values of control epi-NSCs from the same brain; maximal projection of apical Δz = 2 μm) were plotted against apical area (μm^2^) for 102 epi-NSCs (*n* = 5 *optix*>L’sc RNAis brains). An inverse correlation between Neur-GFP intensity and apical area was observed, suggesting that Neur positively regulates apical constriction.

**Supplementary Figure 5:**
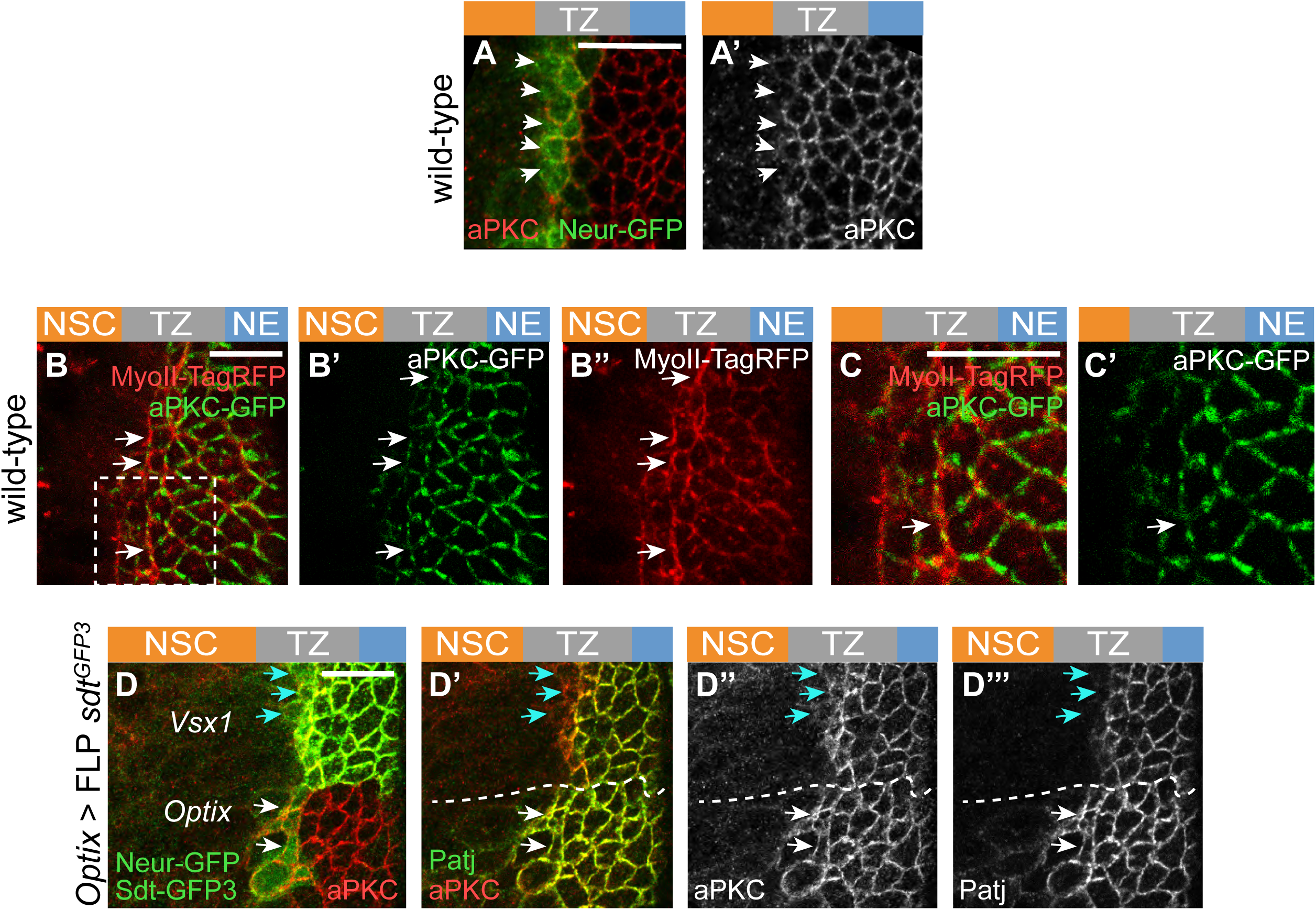
(**A**,**A’**) Low levels of aPKC (red) was detected in epi-NSCs (Neur-GFP, green) in wild-type brains. (**B-C’**) aPKC-GFP (green) localized uniformly at apical junctions of NE and TZ cells in a live brain but appeared to localize in patches at the apical cortex of epi-NSCs. Note that MyoII-TagRFP (red) was observed to be enriched along junctions where aPKC were decreased (white arrows). The inset boxed in B is shown in C,C’. (**D-D’’’**) While low aPKC levels were observed at the apical cortex of control epi-NSCs in the Vsx1 domain (blue arrows), expression of the Neur-resistant isoforms of Sdt upon excision of exon 3 of *sdt* in the Optix domain (hence the loss of Sdt-GFP3, green; arrows) led to the persistent accumulation of aPKC (red; white arrows in L-L’’) and Patj (green; L’,L’’) in epi-NSCs (Neur-GFP, green in L). Scale bars = 10 μm.

**Supplementary Figure 6:**
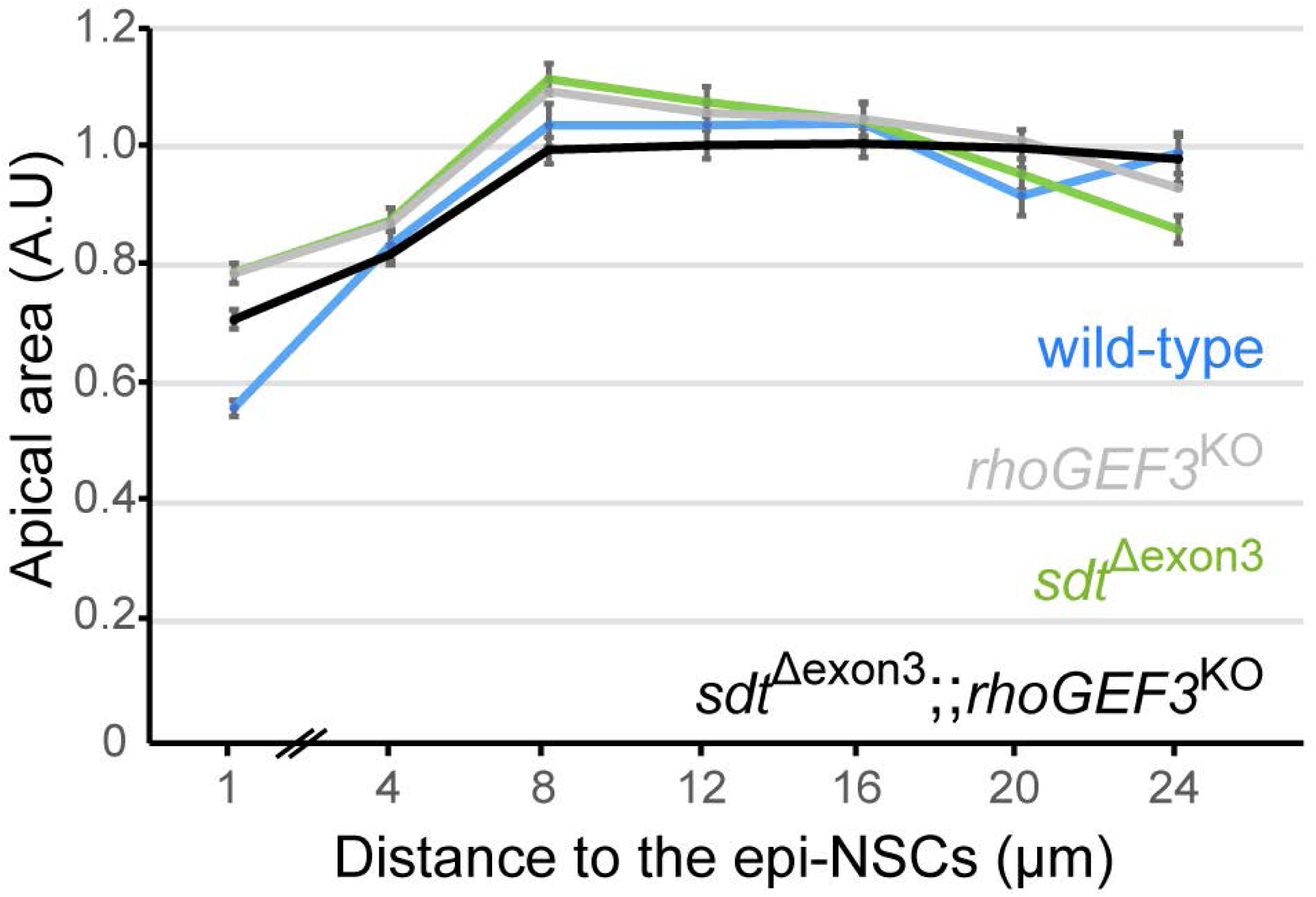
(**A**) Quantification of apical area (normalized, A.U) in the NE of *sdt*^*Δ3*^ *rhoGEF3* double mutants (black; n = 9 brains) plotted against the distance from the TZ medial edge (70-474 cells per binned distance. The phenotype of the double mutant appeared to be similar to those of *rhoGEF3* (data from Fig. 6D) and *sdt*^*Δ3*^ (data from Fig.5E; wild-type data are from Fig. 6D): double mutant cells located within 0-2μm of the TZ medial edge do not have significantly different apical areas than either single mutant (ANOVA).

**Movie S1**

Movie of a brain expressing MyoII-GFP (green) and Par3-mCherry (red) showing an apically constricting TZ cell (outline) undergoing phases of constriction and expansion. Constriction is associated with increased levels of medial-apical MyoII.

**Movie S2**

Movie of a brain expressing MyoII-GFP (green) and Par3-mCherry (red) showing a NE cell (dotted outline) undergoing phases of constriction and expansion. Constriction is associated with increased levels of medial-apical MyoII.

